# Epsin1 enforces a condensation-dependent checkpoint for ubiquitylated cargo during clathrin-mediated endocytosis

**DOI:** 10.1101/2025.02.12.637885

**Authors:** Susovan Sarkar, Hao-Yang Liu, Feng Yuan, Brandon T. Malady, Liping Wang, Jessica Perez, Eileen M. Lafer, Jon M. Huibregtse, Jeanne C. Stachowiak

## Abstract

Clathrin-mediated endocytosis internalizes proteins and lipids from the cell surface, supporting nutrient uptake, signaling, and membrane trafficking. Recent work has demonstrated that a flexible, liquid-like network of initiator proteins is responsible for catalyzing assembly of clathrin-coated vesicles in diverse organisms including yeast, mammals, and plants. How do cells regulate the assembly of this dynamic network to produce cargo-loaded vesicles? Here we reveal the ability of an endocytic adaptor protein, Epsin1, to conditionally stabilize the initiator protein network, creating a cargo-dependent checkpoint during clathrin-mediated endocytosis. Epsin1 is known to recruit ubiquitylated transmembrane proteins to endocytic sites. Using *in vitro* assays, we demonstrate that Epsin1 uses competitive binding and steric repulsion to destabilize condensation of initiator proteins in the absence of ubiquitin. However, when polyubiquitin is present, Epsin1 binds to both ubiquitin and initiator proteins, creating attractive interactions that stabilize condensation. Similarly, in mammalian cells, endocytic dynamics and ligand uptake are disrupted by removal of either ubiquitin or Epsin1. Surprisingly, when Epsin1 and ubiquitin are removed simultaneously, endocytic defects are rescued to near wildtype levels, although endocytic sites lose the ability to distinguish between ubiquitylated and non-ubiquitylated cargos. Taken together, these results suggest that Epsin1 tunes protein condensation to ensure the presence of ubiquitylated cargo during assembly of clathrin-coated vesicles. More broadly, these findings illustrate how a balance of attractive and repulsive molecular interactions controls the stability of liquid-like protein networks, providing dynamic control over key cellular events.

## Introduction

Clathrin-mediated endocytosis (CME) is a fundamental cellular process that internalizes proteins and lipids from the plasma membrane and supports numerous physiological functions including nutrient uptake, signal transduction, synaptic vesicle recycling, and intracellular trafficking^1–3^. CME begins when initiator proteins, including Eps15, Fcho, and Intersectin, assemble on the plasma membrane^2–3^ and subsequently recruit adaptor proteins such as AP2, CALM/AP180, and Epsin1^4,5^. These adaptors then recruit clathrin and transmembrane cargo proteins^5–7^. The adaptor proteins promote the assembly of clathrin into an icosahedral lattice and work together with clathrin to bend the membrane, leading to vesicle budding, scission, and ultimately the formation of clathrin-coated vesicles^5–8^. Notably, CME is a highly stochastic process. As initiators, adaptors, and clathrin triskelia come together at the plasma membrane, some nascent assemblies mature productively into coated vesicles that encapsulate cargo, while others disassemble abortively or stall indefinitely^9–12^. Importantly, initiator proteins, which are some of the first components to arrive at nascent endocytic sites, play a key role in determining which assemblies develop into productive vesicles and which do not^4,13–15^.

Each of the initiator proteins contains substantial regions of intrinsic disorder. These regions contain multiple weak binding motifs for one another such that they facilitate dynamic multivalent binding, a key requirement for biomolecular condensation^16–18^. Recent work has suggested that a flexible network of endocytic initiator proteins is important for the efficient assembly of clathrin-coated vesicles^16^. Specifically, mammalian Eps15 and Fcho1/2, which form biomolecular condensates *in vitro*, assemble into a network which substantially influences the fate of endocytic sites^14^. Stable assembly of this network leads to productive vesicles, while unstable assembly results in abortive structures. Similarly, Ede1, the homolog of Eps15 in budding yeast, forms liquid-like complexes that recruit diverse endocytic adaptor proteins^19^ and the AtEH1/2 subunits of the TPLATE complex, the key endocytic initiator proteins in plants, form liquid-like condensates *in vitro* and in live cells^20^. Together, these studies suggest a conserved propensity of endocytic initiator proteins to assemble into flexible networks that orchestrate the fate of nascent clathrin-coated vesicles.

If the assembly of endocytic initiator proteins controls the fate of endocytic vesicles, how are these assemblies regulated to ensure that productive vesicles contain transmembrane cargo proteins? Among the many types of cargo encapsulated by CME, ubiquitylated transmembrane proteins, marked for recycling and degradation by posttranslational modification, are a major class^21–23^. Eps15 and its homologs use ubiquitin interacting motifs (UIMs) at their C-termini to bind to ubiquitylated proteins^24–27^. These interactions are known to promote assembly of endocytic sites^26^ by promoting condensation of the Eps15 network^15^. However, as an initiator protein, Eps15 is not incorporated into productive coated vesicles. Instead, it remains at the plasma membrane^28,29^, where it catalyzes multiple rounds of endocytosis. Therefore, incorporation of ubiquitylated cargo into clathrin coated vesicles requires adaptor proteins of the Epsin family^8,30–33^. Epsin1 in particular plays a critical role in the internalization of ubiquitylated cargo^34,35^.

Here we asked how Epsin1 works together with Eps15 to ensure the productive assembly of endocytic vesicles that are loaded with ubiquitylated cargo. Our *in vitro* studies revealed that the impact of Epsin1 on condensates of Eps15 is ubiquitin dependent, with Epsin 1 destabilizing condensates in the absence of ubiquitin, yet promoting condensate formation in the presence of polyubiquitin. In mammalian cells, removal of either Epsin1^34,35^ or ubiquitin^15,26^ from endocytic sites disrupted receptor endocytosis, as expected. However, simultaneous removal of both components rescued endocytic dynamics to near wild-type levels, though cargo selectivity was no longer ubiquitin-dependent. Therefore, Epsin 1 enforces a checkpoint for ubiquitylated cargo during clathrin-mediated endocytosis by destabilizing nascent endocytic sites in the absence of ubiquitylated cargo, while promoting maturation of endocytic sites in the presence of ubiquitylated cargo.

## Results

### Epsin1 fails to form protein condensates

Eps15 consists of three N-terminal Eps15 Homology (EH) domains, a central coiled-coil region that facilitates oligomerization, a large intrinsically disordered region (IDR, ∼400 amino acids) which contains multiple DPF (Asp-Pro-Phe) motifs, followed by two UIMs at its C-terminus^36^ (Figure 1a). As previously reported, we found that Eps15 forms liquid-like condensates that fuse together rapidly upon contact (Figure 1b, Supplementary Movie S1). Previous work showed that the coiled-coil domain and interactions between the EH and IDR domains are essential to this condensation^14^. In particular, the coiled-coil domain drives Eps15 dimerization, while EH-IDR interactions likely promote assembly of anti-parallel tetramers^37^. Epsin1 consists of an N-terminal ENTH (Epsin N-Terminal Homology) domain that binds PI(4,5)P2 membrane lipids, followed by two UIMs and an intrinsically disordered C-terminal domain. The disordered region contains three NPF (Asn-Pro-Phe) motifs, which bind EH domain-containing proteins such as Eps15^8^. Given that both proteins have substantial regions of intrinsic disorder, a feature that is often associated with a propensity for condensation^38–41^, we began by asking whether Epsin1, similar to Eps15, formed protein condensates *in vitro*. To assess Epsin1’s potential for condensation, we added Epsin1 at increasing concentrations to a solution containing 3 weight% polyethylene glycol 8000 (PEG8K), which is often used to mimic the crowded cytosolic environment^14,42,43^. We did not observe any evidence of protein condensation for protein concentrations ranging from 20 μM to 80 μM (Figure 1c), substantially above the concentration required for condensation of Eps15 (12 μM). Next, we held the protein concentration at 40 μM and increased the concentration of PEG8K from 3 weight% to 10 weight%, substantially exceeding the concentration required for condensation of Eps15 (3 weight%). Nonetheless, no condensation of Epsin1 was observed (Figure 1d). These experiments suggest that Epsin1 is substantially less prone to condensation compared to Eps15, and that Epsin1 likely resists condensation under physiological conditions. This resistance to condensation may stem from the lack of self-interaction motifs in Epsin1 and electrostatic repulsion between negatively charged amino acids^36,44^. However, when mixed at a 10:1 molar ratio with 16 μM Eps15 (3 weight% PEG8K), Epsin1 (Atto 594 fluorophore) partitioned uniformly into Eps15 condensates (Figure 1e), with a partition coefficient (K_app_) of 8 ± 1, (Figure 1f). To determine if this partitioning was driven by binding between Epsin1’s NPF motifs and Eps15’s EH domains, we created an Epsin1 variant lacking all three NPF motifs (Epsin1ΔNPF), which exhibited significantly weaker partitioning, with a K_app_ of 2 ± 0.3 (Figure 1e-f). As a negative control, amphiphysin, an endocytic protein that contains a long intrinsically disordered region (∼400 amino acids) but no NPF motifs^45^, had a similarly low K_app_ of 2 ± 0.2 (Figure 1f). Importantly, condensates composed of Eps15 and Epsin1 retained their liquid-like properties, fusing and re-rounding upon contact (Figure 1g, Supplementary Movie S2). Collectively, these results suggest that while Epsin1 is unlikely to form condensates on its own, it uses its NFP motifs to interact with condensates composed of Eps15 (Figure 1h).

**Figure 1.**
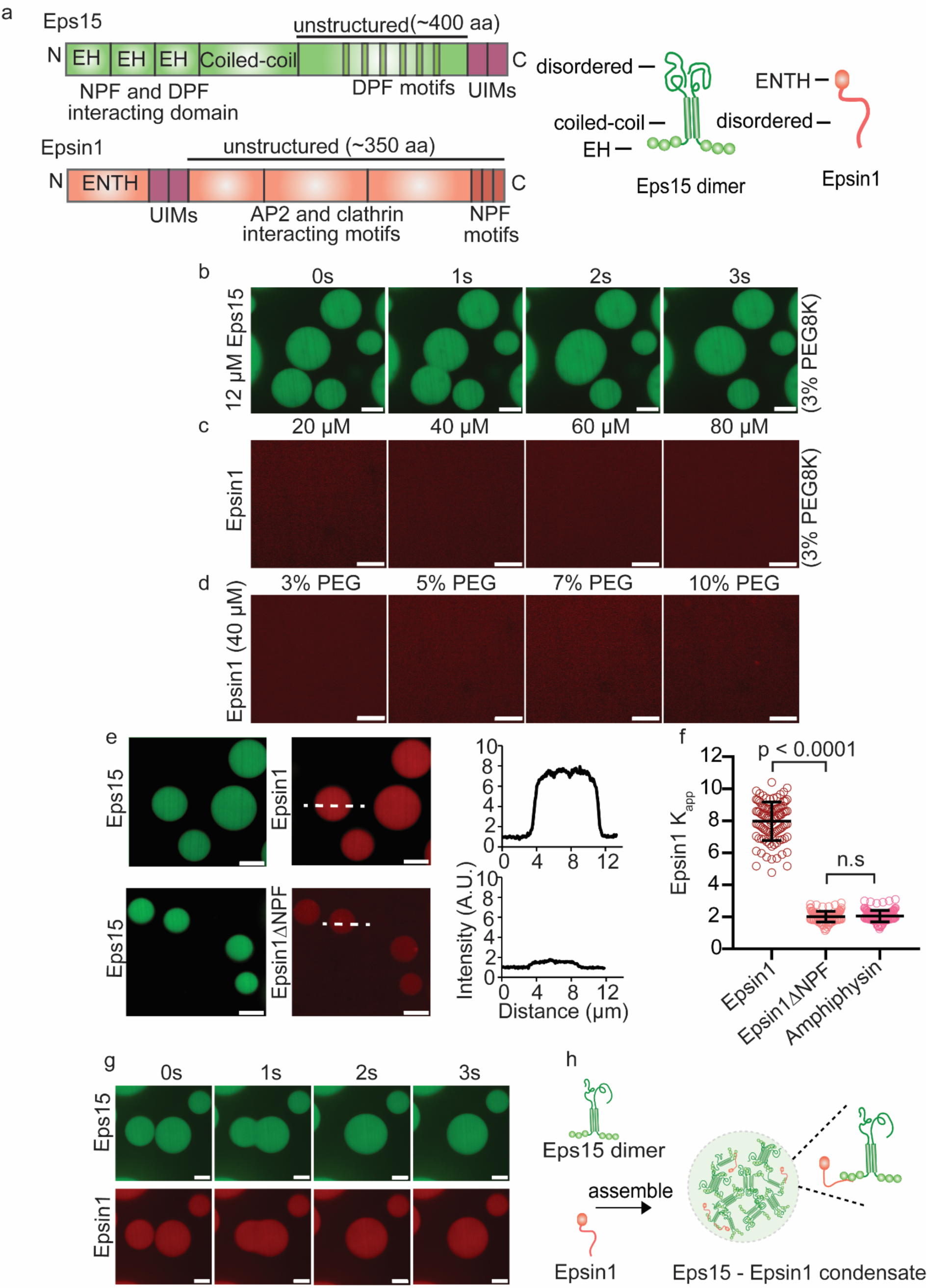
Epsin1 fails to form protein condensates but is incorporated into condensates of Eps15 at low stoichiometric ratios. **a**, Schematic of Eps15 and Epsin1 functional domains. Eps15 consists of three EH domains at its N terminus followed by a coiled-coil domain and a long intrinsically disordered region followed by two ubiquitin interacting motifs (UIMs) at the C terminal end. Epsin1 consists of the N-terminus ENTH (Epsin1 N-Terminal Homology) domain followed by two UIMs. The central part includes the CLAP (Clathrin/AP2 binding) region and multiple DPW motifs for binding AP2 followed by the C-terminus containing 3 NPF (Asn-Pro-Phe) motifs that bind to EH domain-containing proteins like Eps15. The side cartoons depict domain organization of Eps15 in dimeric form and Epsin1 as monomer. **b**, Time course of fusion and re-rounding of Eps15 (12 μM) condensates (green) on contact. **c,** Varying concentration of Epsin1 (20 μM to 80 μM) in 3 weight% PEG8K. **d**, 40 μM Epsin1 in varying weight% of PEG8K (3 weight% to 10 weight%). **e**, Eps15 (16 μM) condensates (green) incubated with 1.6 μM of Epsin1 and Epsin1ΔNPF (red), respectively. Plots on the right depict the intensity profile of the Epsin1 / Epsin1ΔNPF channel along the white dashed line shown in the corresponding images. **f**, The distribution of the Epsin1 and Epsin1ΔNPF intensity ratio (Kapp) between the intensity inside the condensates and the solution. Partition of Amphyphysin was used as a negative control. In total, 100 condensates were analyzed under each condition. Data are mean ± standard deviation, calculated from n = 3 replicates. Statistical significance was tested using an unpaired, two-tailed student’s t test on GraphPad Prism. **g**, Representative time course of fusion events between condensates containing Eps15 and Epsin1. Inset in **h** shows the interaction between Eps15 and Epsin1 (the NPF motif of Epsin1 binds to the EH domains of Eps15). All droplet experiments were performed in 20 mM Tris-HCl, 150 mM NaCl, 5 mM TCEP, 1 mM EDTA and 1 mM EGTA at pH 7.5 with PEG8K, at room temperature, 24 °C. All scale bars equal 5 μm.

### Epsin1 destabilizes condensation of Eps15

We next asked what impact Epsin1 has on the stability of Eps15 condensates. We mixed unlabeled Epsin1 with Eps15 at molar ratios ranging from 8:1 to 1:1. For each ratio, we measured the partitioning of Eps15 between the solution and condensate phases (Figure 2a-c), where higher K_app_ values reflect denser, more stable condensates. We observed a steady decline in Eps15 partitioning to the condensate phase as the ratio decreased from 8:1 to 2:1 (Figure 2a-b). At a ratio of 1:1, condensates no longer formed (Figure 2a). These results suggest that condensation of Eps15 is destabilized by Epsin1 (Figure 2a-b). We also measured the melting temperature (T_m_) for condensates formed in the presence of increasing amounts of Epsin1, where a higher T_m_ indicates a more stable condensate. Condensates consisting of Eps15 alone (16 µM) dissolved at 32°C, in agreement with prior studies (Figure 2c-d)^14,15^. As the concentration of Epsin1 increased, T_m_ steadily decreased, reaching 28°C for a 2:1 mixture of Eps15 and Epsin1 (Figure 2c-e and supplementary Figure S1-S2). These results further confirm that Epsin1 reduces the thermodynamic stability of Eps15 condensates. This destabilizing effect relied on specific interactions between Eps15’s EH domains and Epsin1’s NPF motifs, as Epsin1ΔNPF failed to destabilize condensation of Eps15 when the two proteins were mixed at a 1:1 ratio (Figure 2f). Overall, these results suggest that Epsin1 is capable of destabilizing Eps15 condensates by competing with interactions between Eps15 proteins (Figure 2g).

**Figure 2.**
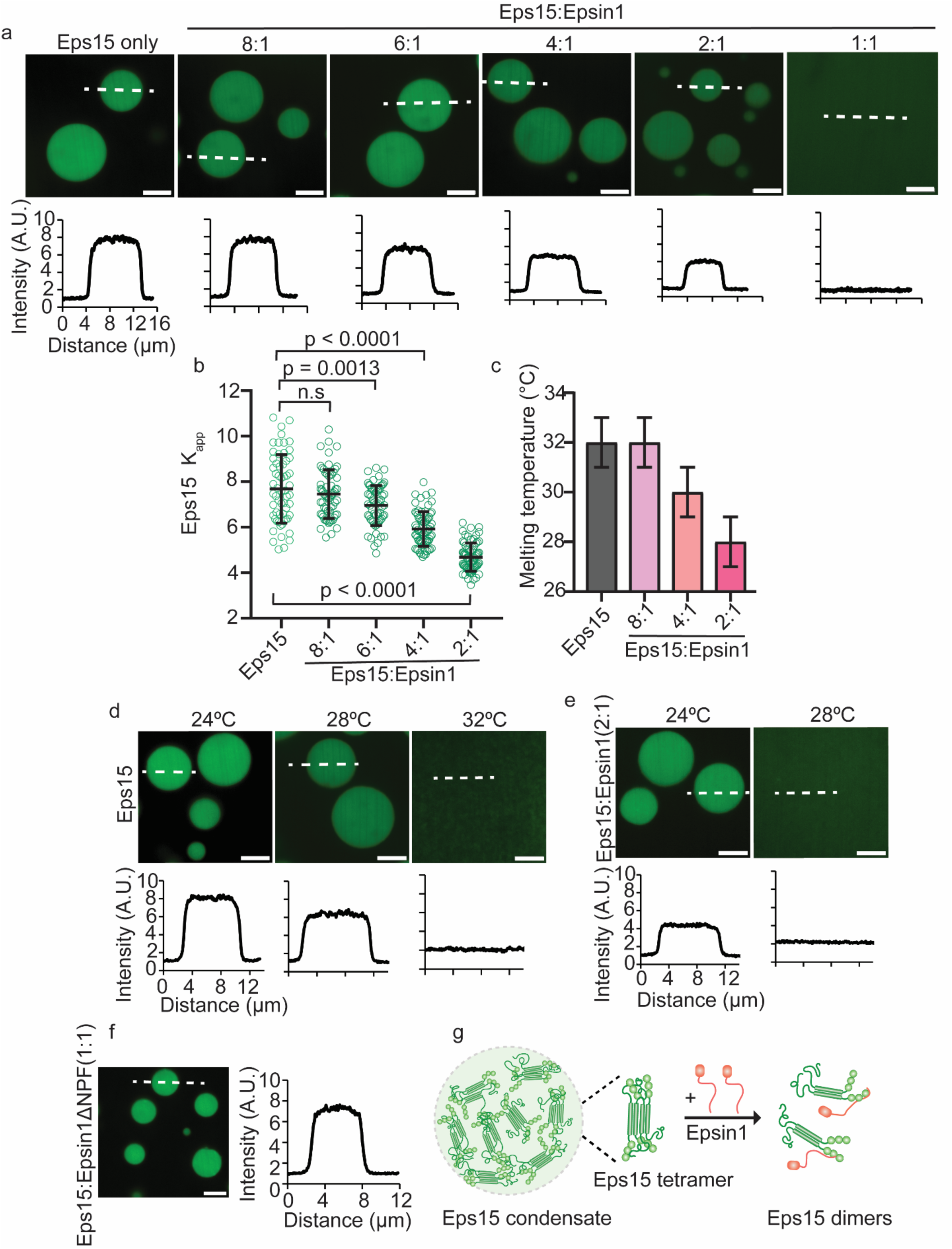
At higher stoichiometric ratios, Epsin1 destabilizes condensation of Eps15. **a,** Representative images of Eps15 condensates with varying stoichiometry of Epsin1. 16 μM Eps15 was used for all the experiments and Epsin1 was mixed with Eps15 from 8:1 to 1:1 molar ratio. The line profiles depict the intensity profile of green (Eps15) channels along the white dashed line. **b,** The distribution of the Eps15 intensity ratio (Kapp) between the intensity inside the condensates and the solution with varying molar ratio of Epsin1. In total 100 condensates were analyzed under each condition. Data are mean ± standard deviation, calculated from n = 3 replicates. Statistical significance was tested using an unpaired, two-tailed student’s t test on GraphPad Prism. **c,** Bar graph shows the melting temperature (Tm) of Eps15 and Eps15:Epsin1 mixed condensates, n = 3 replicates. The error bar represents ± 1°C of the melting temperature. **d-e,** Representative images of protein condensates at increasing temperatures. Plots show fluorescence intensity of Eps15 measured along dotted lines in each image. Condensates are formed from **(d)** 16 μM Eps15, **(e)** 16 μM Eps15 with 8 μM Epsin1. **f,** Eps15 (16 μM) condensates mixed with 16 μM (1:1) of Epsin1ΔNPF, the line profile shows the intensity of Eps15 across the white dashed line. **g,** Shows the cartoon of the Eps15 multimerization through intermolecular interaction between DPF motif and EH domains and how the addition of Epsin1 can destabilize Eps15 condensates. Epsin1 binds to the EH domain of Eps15 and destabilizes the condensates by preventing Eps15 multimerization. All condensate experiments were performed in 20 mM Tris-HCl, 150 mM NaCl, 5 mM TCEP, 1 mM EDTA and 1 mM EGTA at pH 7.5 with PEG8K, at room temperature, 24 °C. All scale bars equal 5 μm.

### When ubiquitin is present, Epsin1 stabilizes condensation of Eps15

Given that Eps15, Epsin1, and ubiquitin are all present simultaneously at endocytic sites^26,27,34,35^, we next investigated the impact of ubiquitin on the tendency of Eps15 and Epsin1 to form condensates. It was previously established that poly-ubiquitin, but not mono-ubiquitin, partitions favorably to Eps15 condensates and stabilizes their assembly^15^. Therefore, we first confirmed that K63-linked tetra ubiquitin (TetraUb), partitioned positively into liquid-like condensates composed of Eps15 and Epsin1 (Figure 3a and Supplementary Movie S3). Next, we investigated the impact of Epsin1 on this partitioning. Specifically, we mixed Eps15 and Epsin1 in ratios from 8:1 to 2:1, and measured the partitioning of TetraUb (Atto 646 labeled) into the resulting condensates (Figure 3b,c). Interestingly, addition of a small amount of Epsin1 to condensates composed of Eps15 (8:1 Eps15:Epsin1) increased the partitioning of TetraUb, suggesting that polyubiquitin may cross-link UIM domains within Eps15 and Epsin1 proteins. However, further increasing the relative concentration of Epsin1 (6:1 to 2:1 Eps15:Epsin1) led to a progressive decrease in TetraUb partitioning, likely owing to destabilization of the Eps15 network by increasing amounts of Epsin1 (Figure 2a-c). Similarly, the presence of TetraUb, but not mono ubiquitin (MonoUb), resulted in stronger partitioning of Epsin1 to condensates (10:1 Eps15 and Epsin1), an effect which required the UIM domains of both Eps15 and Epsin1 (Supplementary Figure S3). Further, while we showed in Figure 2a that a 1:1 solution of Eps15 and Epsin1 does not form condensates, adding TetraUb at increasing concentrations to this solution caused protein condensates to progressively reemerge (Figure 4d-e). Together, these results suggest that when polyubiquitin is present, Epsin1 loses its ability to destabilize protein condensates and can instead stabilize them. To test this idea, we measured the impact of Epsin1 (8:1 Eps15:Epsin1) on the melting temperature (T_m_) of protein condensates in the presence and absence of TetraUb, Figure 3f, (Supplementary Figure 5). First, we established that condensates composed of Eps15 and Epsin1 in an 8:1 ratio had a similar melting temperature (32°C) to condensates composed purely of Eps15 (Supplementary Figure S1, S4), suggesting that small amounts of Epsin1 do not substantially destabilize Eps15 condensates. Previous work demonstrated that polyubiquitin stabilizes Eps15 condensates by cross-linking Eps15’s UIMs^15^. In line with these results, we found that addition of 0.12 µM TetraUb increased the T_m_ of Eps15 condensates to 36°C in the absence of Epsin1 (Figure 3f, supplementary Figure S6). However, addition of TetraUb at the same concentration to condensates consisting of both Eps15 and Epsin1 (8:1) led to an even higher T_m_ of 38°C, (Figure 3f, Supplementary Figure S4-S6). Collectively, these results suggest that, while Epsin1 destabilizes condensation of Eps15 alone (Figure 2), in the presence of polyubiquitin, Epsin1 is capable of stabilizing protein condensates (Figure 3g).

**Figure 3:**
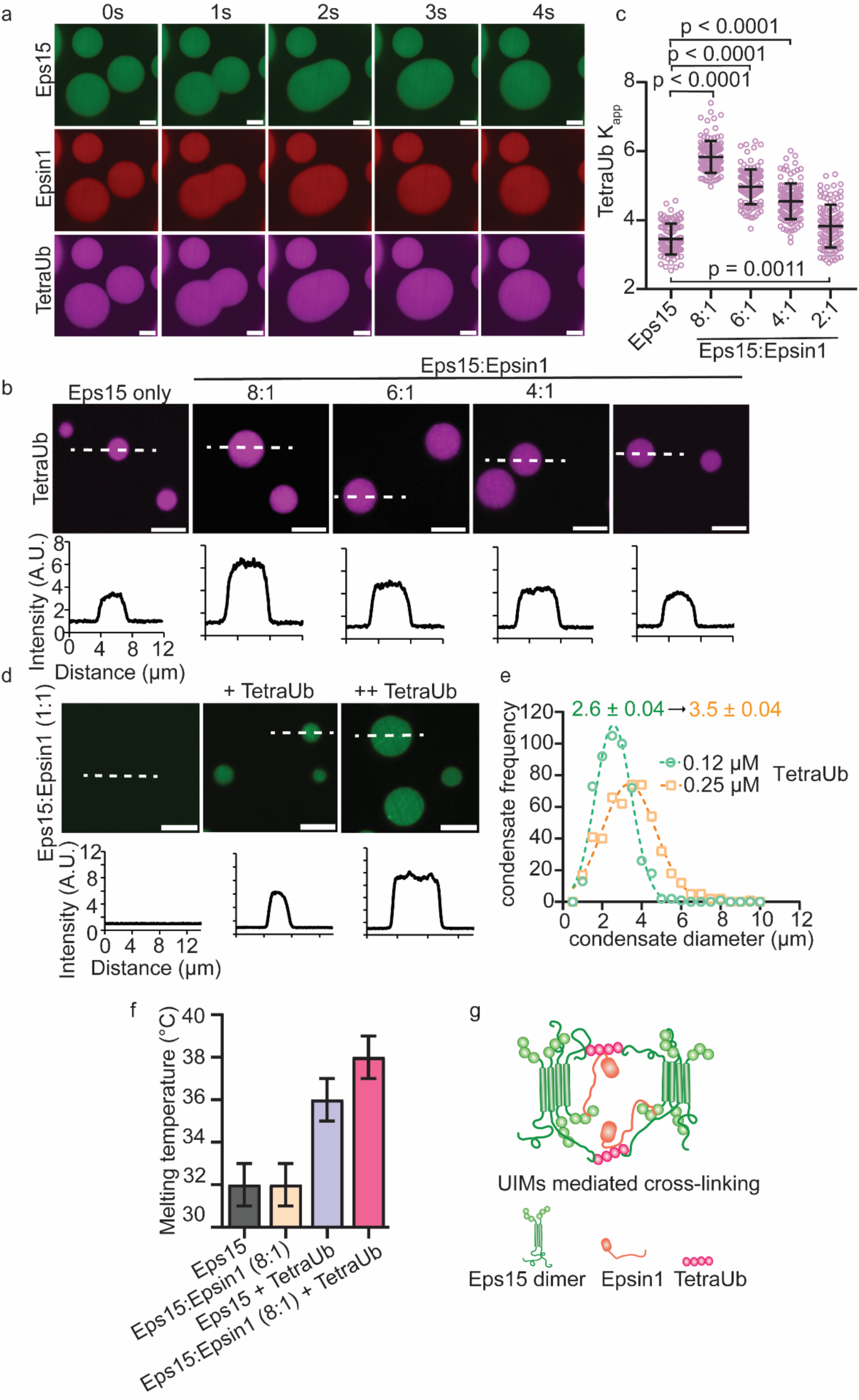
**a,** Representative time course of fusion events between Eps15 (16 μM) condensates (green) with 1.6 μM of Epsin1 (red) and 0.1 μM K63 linkage TetraUb (magenta), respectively. **b,** Representative images of Eps15 condensates with varying molar ratio of Epsin1 (8:1 to 2:1) in presence of TetraUb (magenta). The corresponding line profiles depict the intensity profile of the TetraUb (magenta) channel along the white dashed line. **c,** The distribution of the TetraUb (magenta) intensity ratio (Kapp) between the intensity inside the condensates and the solution with varying molar ratio of Epsin1. In total 100 condensates were analyzed under each condition. Data are mean ± standard deviation, calculated from n = 3 replicates. Statistical significance was tested using an unpaired, two-tailed student’s t test on GraphPad Prism. **d,** Representative images of the condensates containing 16 µM of Eps15 (green) and 16 µM of Epsin1 (1:1 ratio) in addition to 0.12 µM and 0.25 µM of TetraUb, respectively. The corresponding line profile depicts intensity profile of green (Eps15) channel along the white dashed line, n = 3. **e,** The size distribution histogram in Gaussian fit of the condensates containing 16 µM of Eps15 and Epsin1 in presence of different (0.12 and 0.25 µM) TetraUb concentration. In total 200 condensates were analyzed under each condition from n = 3 replicates. **f,** The bar plot depicts the melting temperature (Tm) of condensates composed of Eps15, Eps15 mixed with Epsin1 (8:1), Eps15 with TetraUb and when all three present together, n = 3 replicates. The error bar represents ± 1°C of the melting temperature. **f,** Shows the cartoon of UIM mediated crosslinking between Eps15 and Epsin1 that can stabilize the condensates. All condensate experiments were performed in 20 mM Tris-HCl, 150 mM NaCl, 5 mM TCEP, 1 mM EDTA and 1 mM EGTA at pH 7.5 with PEG8K, at room temperature, 24°C. All scale bars equal 5 μm.

**Figure 4.**
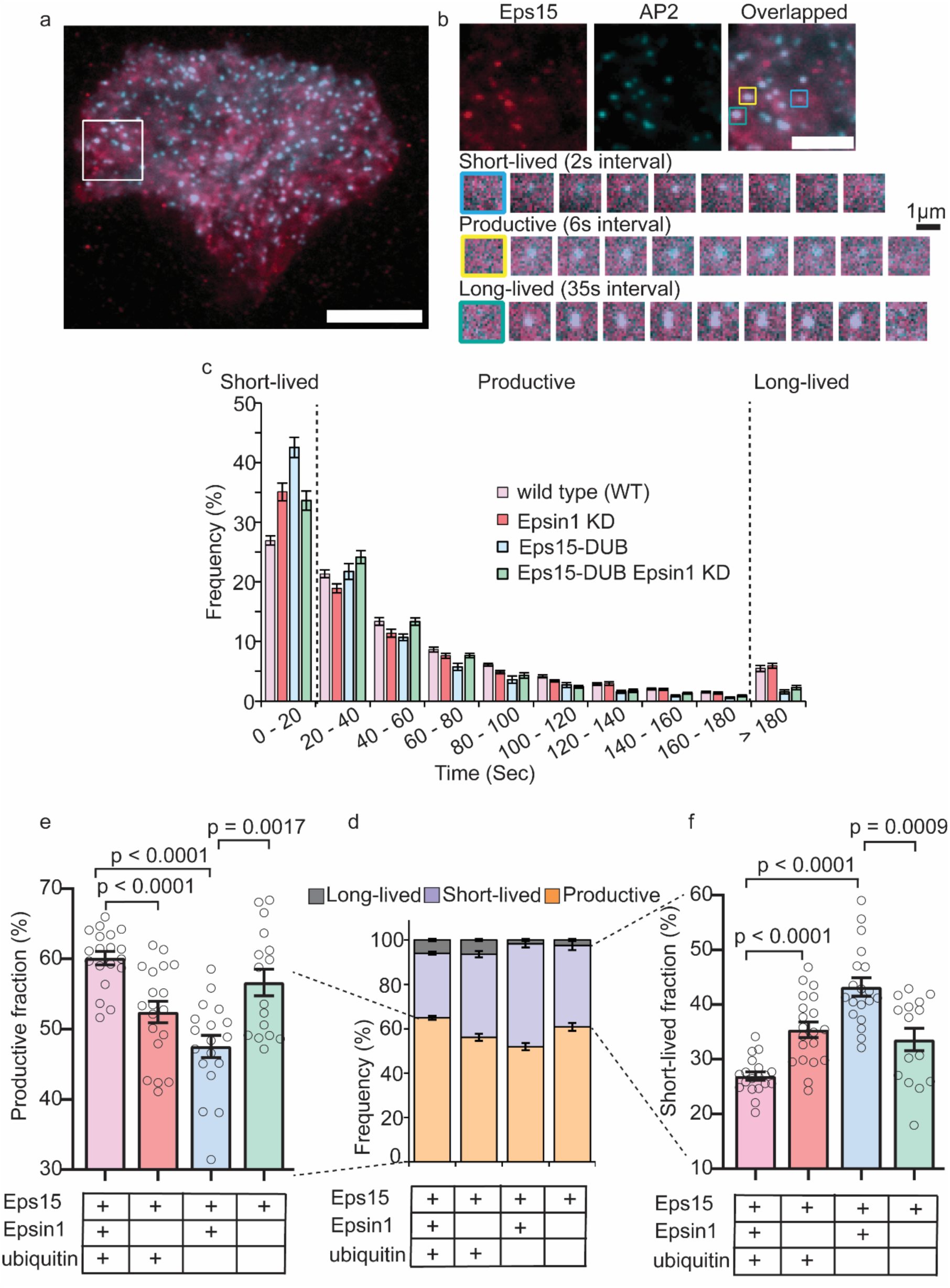
Loss of ubiquitin from endocytic sites rescues defects in endocytic dynamics resulting from knock down of Epsin1. **a,** Representative image of a SUM cell expressing gene-edited AP2 σ2-HaloTag: JF646 (cyan) and Eps15-mCherry (red). The scale bars 10 μm. Large inset highlights three representative clathrin-coated structures shown in smaller insets. The scale bars 5 μm. **b**, Representative image of short-lived (blue), productive (yellow) and long-lived (green) structures lasting < 20 s, > 20 s and > 5 min, respectively. **c,** Histograms of lifetime distributions of clathrin-coated structures under different experimental groups. Lifetime shorter than 20 s is considered short-lived, lifetime between 20 and 180 s is labeled as productive and structures lasting longer than 180 s are long-lived. Data are mean ± S.E.M. WT represents SUM cells that have endogenous Eps15 and Epsin1 expression. Eps15 KO represents SUM cells that were CRISPR modified to knock out alleles of endogenous Eps15. Epsin1 KD represents Epsin1 knock down in WT cells using siRNA. Eps15-DUB represents Eps15 knock out cells transfected with Eps15 fused with DUB (to C terminal end of Eps15). Eps15-DUB Epsin1 KD represents Eps15 knock out cells transfected with Eps15 fused with DUB and knocked down for Epsin1. All three constructs have mCherry at their C terminus for visualization. **d,** Bar plot summarizing percentage of short-lived (abortive), productive and long-lived fraction for the four conditions. **e,** Bar plot of the productive fraction for all four groups. Data are mean ± S.E.M. **f,** Bar plot of the short-lived fraction for all four groups. Data are mean ± S.E.M. For WT, Epsin1 KD, Eps-DUB and Eps-DUB Epsin1 KD, n = 20, 20, 15 and 16 biologically independent cells, respectively, were imaged. In total > 10000 pits were analyzed for each group. An unpaired, two-tailed student’s t test was used for statistical significance using GraphPad prism, n.s. means no significant difference. p < 0.05 is considered significantly different. All cell images were collected at 37°C.

### Epsin1 and ubiquitylation play compensatory roles in endocytic dynamics

We next sought to determine how interactions between Epsin1 and ubiquitin influence the dynamics of endocytic events in living cells. Prior research has established ubiquitin as a crucial stabilizer of the early endocytic protein complex, of which Eps15 is a major component^15^. The assembly of endocytic structures at the plasma membrane is highly stochastic and dynamic^46,47^. Early in the process, endocytic initiator proteins, including Eps15, arrive at the plasma membrane, where they subsequently recruit multiple endocytic adaptor proteins, including Epsin1 and AP2, which in turn recruit clathrin^1–3^. The majority of these nascent assemblies mature into productive endocytic vesicles^47^. However, a substantial minority, approximately 30%, disassemble prematurely leading to “short-lived” endocytic events that typically reside at the plasma membrane less than 20 seconds and fail to internalize cargo^47^. Conversely, productive assemblies, which result in vesicle formation and cargo internalization, remain at the plasma membrane for periods ranging from 20 seconds to several minutes^41^. At the opposite end of the spectrum, endocytic structures that persist for longer than a few minutes are considered “long-lived”^47,48^. These overly stable assemblies are often larger than productive endocytic structures and may be removed from the plasma membrane by autophagy rather than endocytosis^49^. Shifts in the dynamics of endocytic structures can serve as an indicator of endocytic efficiency^14,46–48^. An increase in short-lived structures suggests reduced stability, while an increase in productive or long-lived structures indicates enhanced stability. We examined the impact of Eps15, Epsin1, and ubiquitin on the distribution of short-lived, productive, and long-lived endocytic structures in mammalian cells.

Our studies employed a human breast cancer-derived epithelial cell line (SUM159), which was gene-edited to fuse a halo-tag to the C-terminus of the σ2 subunit of adaptor protein 2 (AP2)^50^ for the purpose of visualizing the assembly and dynamics of endocytic sites. A second cell line was created by further gene editing these cells to knock out Eps15 for the purpose of replacing it with modified variants, as described below.^14^ These cell lines will be henceforth referred to as wild type cells and Eps15 knock out (Eps15 KO) cells, respectively. We have previously established that defects in endocytic dynamics resulting from deletion of Eps15 are fully rescued by transient expression of a version of Eps15 with an mCherry tag fused to its C-terminus^14,15^. AP2 is a critical adaptor protein that is thought to be present at endocytic sites in a nearly 1:1 stoichiometric ratio with clathrin^51^. Fluorescent tagging of AP2 (JF646 bound to halotag^52^) allowed us to visualize and track endocytic structures in real-time during live-cell imaging. Notably, it is preferable to track AP2 rather than clathrin because AP2 localizes exclusively to the plasma membrane^51^. We conducted live cell imaging experiments using total internal reflection fluorescence (TIRF) microscopy to quantify endocytic dynamics. In TIRF images, endocytic structures appeared as diffraction-limited fluorescent puncta in the AP2 channel (JF646, Figure 4a). Based on established classifications^46,48,53^ we defined endocytic structures with lifetimes shorter than 20 seconds as “short-lived”, those lasting between 20 and 180 seconds as “productive,” and those persisting longer than 180 seconds as “long-lived” (Figure 4b).

In wild type cells, we observed endocytic structures falling into each of these categories (Figure 4c-d). We first compared the dynamics of endocytic structures in wild type cells to those in wild type cells in which Epsin1 was knocked down using siRNA (supplementary Figure S7). Consistent with previous reports^54^, the fraction of productive structures decreased significantly (60 ± 4% to 52 ± 6%) upon Epsin1 knock down, mirrored by an increase in short-lived structures (27 ± 3 % to 36 ± 6%), while the long-lived fraction changed only slightly (5.7 ± 2 to 6 ± 2); Figure 4e. Next, to probe the effect of removal of ubiquitin from endocytic sites, we transiently expressed a version of Eps15 linked to a deubiquitylating enzyme (DUB) at its C-terminus (Eps15-DUB) in Eps15 KO cells. Here we used a “broad-spectrum” DUB, UL36, which is expected to disassemble both K63 and K48-linked polyubiquitin chains^55–57^. In a recent study, expression of Eps15-DUB resulted in a significant increase in abortive endocytic structures^15^, highlighting the role of ubiquitin in the assembly of clathrin-coated vesicles. Similarly, in our experiments, the expression of Eps15-DUB resulted in a significant decrease in the fraction of productive (47 ± 6%) and long-lived (1.5 ± 1%) endocytic events, coupled with an increase in the fraction of short-lived endocytic events (43 ± 7%).

We next asked what happens when ubiquitin and Epsin1 are simultaneously removed from endocytic sites. Because Epsin1^32,34,35,54^ and ubiquitin^15,24,26,58^ have each been shown to play important roles in endocytosis, the conventional expectation would be that loss of both components simultaneously would lead to an even larger endocytic defect than loss of either component individually. However, our *in vitro* results showed that Epsin1 destabilizes assembly of Eps15 in the absence of polyubiquitin (Figure 2), such that removal of Epsin1 in the absence of ubiquitin might be expected to increase the stability of nascent endocytic sites. Interestingly, upon simultaneously removing Epsin1 and ubiquitin (Epsin1 knock down and transient expression of Eps15-DUB in Eps15 KO cells), endocytic dynamics were largely rescued, such that the fraction of productive endocytic structures increased relative to the single knock out experiments (56 ± 7%), while the fraction of short-lived structures decreased (33 ± 8%; Figure 4e-f). In control experiments, we observed that scrambled siRNA had no impact on endocytic dynamics and that transient expression of Eps15 fused to a catalytically inactive DUB (Eps15-DUB-dead) rescued the Eps15 knock out phenotype, similar to transient expression of wild type Eps15 (Supplementary Figures S8-S9). Taken together, our results suggest that Epsin1 serves as a checkpoint for the presence of ubiquitin at nascent endocytic sites. When ubiquitin is present, it overcomes the inhibitory effect of Epsin1 on Eps15, leading to productive endocytosis, while an absence of ubiquitin results in Epsin1-dependent destabilization of nascent sites.

### Loss of Epsin1 and ubiquitylation from endocytic sites rescues transferrin uptake

Having established that simultaneous removal of Epsin1 and ubiquitin from endocytic sites rescues endocytic dynamics (Figure 4e-f), we next evaluated the impact of these modifications on endocytic recycling. Specifically, we used flow cytometry to quantify uptake of fluorescent-labeled transferrin (Atto 488) by the cell’s endogenous population of transferrin receptors (TfRs, Figure 5a). Owing to constitutive uptake of TfR by the clathrin pathway via binding interactions with the adaptor protein, AP2^59–61^, transferrin uptake has frequently been used to evaluate the overall productivity of CME^62^. Cells were exposed to labeled transferrin for 30 minutes, followed by trypsinization and washing to remove unbound and surface bound ligands. In wild type cells we observed a substantial fluorescence shift upon exposure to transferrin (compare Figure 5b-c to Supplementary Figure S10), indicating uptake. Epsin1 knock down led to a significant decrease in transferrin uptake (Figure 5b-c and Supplementary Figures S11-S12), as expected^63,64^. Removal of ubiquitin from endocytic sites via transient expression of Eps15-DUB also impaired transferrin uptake, consistent with previous work^15^. In agreement with our data on endocytic dynamics (Figure 4), simultaneous removal of Epsin1 and ubiquitin (Epsin1 knock down and transient expression of Eps15-DUB) restored transferrin uptake to levels similar to wild type cells (Figure 5b-c). Taken together, these seemingly counterintuitive results are consistent with the idea that Epsin1 acts as a checkpoint for ubiquitylated cargo during endocytosis. Specifically, ubiquitin reinforces endocytic assemblies, driving endocytosis forward. In contrast, when ubiquitin is removed from endocytic sites, Epsin1 destabilizes assembly of the Eps15 network, reducing the probability that nascent endocytic sites will mature into productive vesicles (Figure 4e-f and Figure 5c, e). This switch-like mechanism is consistent with the findings of our *in vitro* experiments, which showed that Epsin1 changes from dissolving protein condensates in the absence of ubiquitin (Figure 2) to reinforcing them in its presence (Figure 3). Interestingly, while removal of Epsin1 creates an endocytic defect (Figure 4e-f and 5c), when Epsin1 and ubiquitylation are absent simultaneously, endocytic function and transferrin uptake are rescued (Figure 4e-f and 5c), consistent with the counterbalancing effects of Epsin1 and ubiquitin on Eps15 condensates *in vitro* (Figure 2a-c and 3d). From a cellular perspective, this result suggested that in the absence of both Epsin1 and ubiquitylation, endocytic sites prioritize other cargos and adaptors, which continue to drive endocytosis forward. Therefore, we next asked how the simultaneous loss of ubiquitin and Epsin1 impacts the ability of endocytic sites to detect and prioritize ubiquitylated cargo.

**Figure 5.**
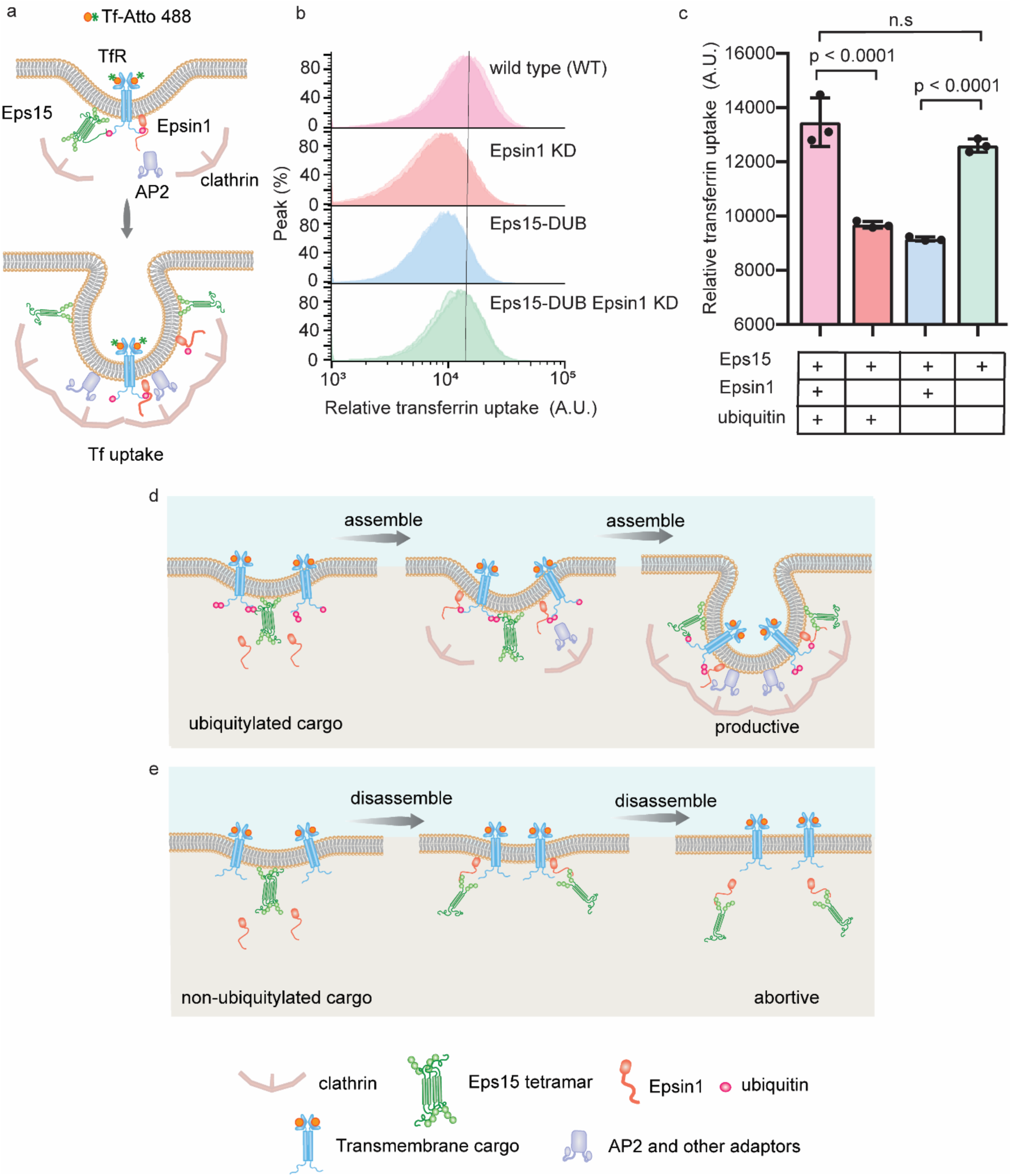
Loss of ubiquitin from endocytic sites rescues defects in transferrin uptake resulting from knock down of Epsin1. **a,** Illustration showing TfR uptake at the plasma membrane. **b,** Flow cytometry histogram of the transferrin fluorescence intensity (green fluorescence) of cells in each condition. **c,** Bar graph represents transferrin fluorescence intensity measured by flow cytometry. Data are mean ± standard deviation, calculated from n = 3 biological replicates. WT represents SUM cells that have endogenous Eps15 and Epsin1 expression. Eps15 KO represents SUM cells that were CRISPR modified to knock out alleles of endogenous Eps15. Epsin1 KD represents Epsin1 knock down in WT cells using siRNA. Eps15-DUB represent Eps15 knock out cells transfected with wild type Eps15 fused with DUB (deubiquitylase fused to C terminal end of Eps15). Eps15-DUB Epsin1 KD represents Eps15 knock out cells transfected with wild type Eps15 fused with DUB and knocked down for Epsin1. All three constructs have mCherry at their C terminus for visualization. An unpaired, two tailed student’s t test was used for statistical significance using GraphPad Prism. P < 0.05 is considered significantly different. Flow cytometry runs were carried out at 24°C. **d-e** Schematic showing how polyubiquitin stabilizes the endocytic protein network by interacting with Eps15 and Epsin1 and how Epsin1 functions as a checkpoint for ubiquitylated cargo. **(d)** In presence of polyubiquitin Eps15 and Epsin1 forms a liquid-like initiator network resulting in productive clathrin-mediated endocytosis. **(e)** In absence of ubiquitylated cargo Epsin1 interacts with Eps15 and prevents the assembly of the liquid-like network of Eps15 and thereby acting as a checkpoint for ubiquitylated cargo to prevent unproductive endocytic events.

### Loss of Epsin1 and ubiquitin from endocytic sites disrupts selective internalization of ubiquitylated cargo

How can CME accommodate the loss of both Epsin1 and ubiquitin while maintaining near wild type dynamics and transferrin uptake? Surely some loss of function must result from the removal of these two important components from endocytic sites. If Epsin1 functions as a checkpoint for ubiquitylated cargo, as we have hypothesized, then we might expect its removal to result in endocytic sites losing their ability to selectively incorporate such cargo. To test this hypothesis, we compared the recruitment of permanently ubiquitylated and non-ubiquitylated model transmembrane proteins into endocytic sites of wild type cells and cells for which endocytic sites lacked both Epsin1 and ubiquitin (Eps15-DUB Epsin1 knock down). The non-ubiquitylated model protein (0-Ub) consisted of the intracellular and transmembrane domains of the transferrin receptor, followed by an extracellular GFP domain. The ubiquitylated model protein (1-Ub) was identical to its non-ubiquitylated counterpart except for the fusion of one ubiquitin to its intracellular N-terminus (Figure 6a). To prevent endogenous ubiquitylation and deubiquitylation by cellular enzymes, we mutated all lysine residues in the intracellular domain and ubiquitin domain to arginine and replaced the C-terminal Gly-Gly residues of ubiquitin with Ala-Ala, respectively. Here we chose a mono-ubiquitylated model cargo, rather than a poly-ubiquitylated version, because we wanted to measure the ability of endocytic sites to sense ubiquitylated cargo, without further cross-linking the endocytic network, as might occur if a poly-ubiquitylated cargo were introduced.

**Figure 6:**
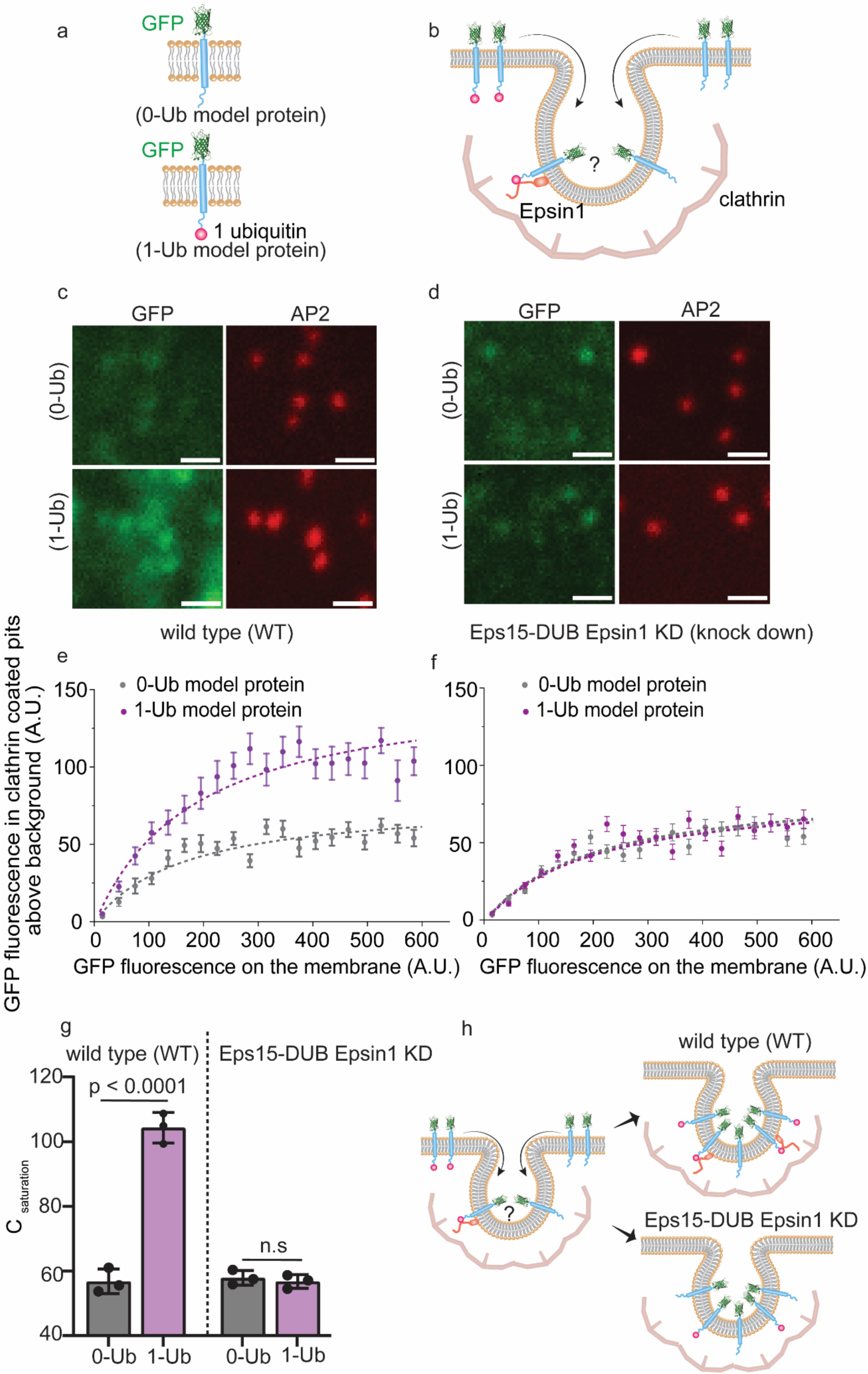
The loss of Epsin1 and ubiquitin impairs cargo selectivity but maintains bulk endocytosis. **a,** Schematic representation of model proteins with defined ubiquitin modifications; the 0-Ub model protein with no attached ubiquitin and the 1-Ub model protein with a single attached ubiquitin. All lysine residues in the cytoplasmic domain were mutated to arginines to prevent ubiquitylation. To further stabilize the attached ubiquitin moieties, all lysines within ubiquitin were mutated to arginines, and the C-terminal GG residues were altered to prevent intracellular deubiquitylation. **b,** Shows the illustration and recruitment of ubiquitylated and non-ubiquitylated model cargo proteins in the endocytic structure. **c-d,** spinning disk confocal images of the plasma membrane of cells transiently expressing the 0-Ub and 1-Ub model proteins. Green fluorescence (GFP) highlights the model proteins, while red fluorescence (AP2-JF647) marks endocytic structures. The scale bars are 500 nm (insets). **e-f,** The relative number of model proteins localized in clathrin-coated structures is shown against the relative concentration of model proteins on the plasma membrane around each structure. Data were collected from over 10000 endocytic sites in n = 30 independent cells, cells expressing 0-Ub and 1-Ub model protein. Each point reflects the average value from clathrin-coated structures binned by the local membrane concentration of the proteins. Data are mean ± standard deviation, while dashed lines represent model predictions based on the best-fit values. **g,** Bar plot shows the relative Csaturation (maximum local concentration of fusion proteins in clathrin-coated structures) values for the 0-Ub and 1-Ub in WT and Eps15-DUB Epsin1 KD cells. Each bar displays the mean ± standard deviation. An unpaired, two tailed student’s t test was used for statistical significance using GraphPad Prism. p < 0.05 is considered significantly different. **h,** Shows the schematic illustration of outcomes of cargo sorting at endocytic sites, in WT cells, ubiquitylated cargo (1-Ub) gets selectively enriched at the site over non-ubiquitylated cargo; in Eps15-DUB Epsin1 KD cells, the selective enrichment of 1-Ub at the endocytic site is lost.

Confocal imaging revealed significant colocalization of both 0-Ub and 1-Ub proteins with endocytic structures in both groups of cells, consistent with the known interaction of the intracellular domain of TfR with AP2^60,61^ (Figure 6c-d and Supplementary Figure S13). However, in wild type cells, the 1-Ub protein appeared to colocalize more strongly with endocytic sites in comparison to the 0-Ub protein (Figure 6c), while 1-Ub and 0-Ub appeared to colocalize equally with endocytic sites in Eps15-DUB, Epsin1 knock down cells (Figure 6d). Using CMEAnalysis^9^, we quantified the partitioning of these model proteins at endocytic sites. The algorithm identified clathrin-coated structures (diffraction limited puncta in AP2 channel) and measured the intensity of model proteins (GFP channel) colocalized with them, relative to their intensity on the surrounding plasma membrane^9,65^. In wild type cells, both 0-Ub and 1-Ub proteins showed an initial increase in recruitment to endocytic sites with increasing plasma membrane concentration, followed by saturation at higher concentrations (Figure 6e). The saturation value for 1-Ub (104 ± 4) was nearly double that of 0-Ub (57 ± 4), indicating that endocytic sites in wild type cells selectively concentrated ubiquitylated cargo (Figure 6e-g). In contrast, experiments in Eps15-DUB Epsin1 knock down cells revealed no significant difference in saturation value between 0-Ub and 1-Ub proteins (Figure 6f-g), suggesting an inability of endocytic sites in these cells to distinguish between ubiquitylated and non-ubiquitylated cargo. Notably, the 0-Ub protein was internalized about equally by both wild type and Eps15-DUB Epsin1 knock down cells, further confirming that simultaneous removal of Epsin1 and ubiquitin from endocytic sites rescues endocytic uptake of non-ubiquitylated transmembrane proteins. Taken together, these results suggest that, while endocytic dynamics (Figure 4) and receptor recycling (Figure 5) are largely normal when ubiquitin and Epsin1 are removed from endocytic sites, the loss of these components renders cells incapable of prioritizing ubiquitylated transmembrane proteins for internalization.

## Discussion

Significant efforts have focused on elucidating the key requirements or “checkpoints” which govern which endocytic assemblies mature into productive clathrin-coated vesicles and which do not ^2,10,12,53,66^. Mettlen, et al., reported checkpoints that monitor clathrin-coated pit assembly, membrane curvature, and the presence of cargo, suggesting that abortive endocytic structures are indicative of a fidelity-monitoring process^12^. Conversely, Pedersen, et al., challenged the notion that cargo primarily influences the initiation of endocytic sites, proposing instead a regulatory transition point that governs the shift from the initiation phase to the internalization phase^10^. This transition was thought to be influenced by cargo, which accelerated the maturation, rather than initiation, of endocytic sites^10^. In both models, checkpoints are thought to be regulated by complex interactions among endocytic proteins, adaptors, and cargo, where post translational modifications are hypothesized to play a significant role^2,10,12,15,53^. However, the specific mechanisms and molecular interactions that encode these checkpoints have remained elusive.

In this study, we demonstrate that Epsin1 functions as a checkpoint for ubiquitylated cargo. Specifically, our results suggest that Epsin1 acts as a biomolecular sensor, detecting ubiquitin as a cargo marker, and thereby regulating the formation of productive endocytic structures in a condensation-dependent manner. *In vitro* experiments showed that Epsin1 does not form condensates independently but integrates into Eps15 condensates through its NPF motifs (Figure 1c-g). In the absence of ubiquitin, Epsin1 destabilizes Eps15 condensates (Figure 2a-e). However, addition of poly-ubiquitin promotes multivalent interactions between Eps15 and Epsin, such that Epsin1 switches from destabilizing protein condensation to stabilizing it (Figure 3).

Cell dynamics and transferrin uptake assays revealed that removing Epsin1, or ubiquitin separately leads to defects in productive endocytosis (Figure 4d-f), consistent with previous reports^14,15,54^. Remarkably, however, simultaneous removal of both Epsin1 and ubiquitin rescued endocytic dynamics and transferrin uptake to near wild type levels (Figure 4d-f and Figure 5b-c). This counterintuitive finding is consistent with our *in vitro* observation that Epsin1 and ubiquitin have opposite effects on the stability of Eps15 condensates. Nonetheless, removal of Epsin1 and ubiquitin was not without effect, as it resulted in endocytic sites losing the ability to selectively concentrate ubiquitylated cargo (Figure 6), a key requirement for maintaining the integrity of the plasma membrane^25,67,68^. Together these findings suggest that Epsin1 serves as a biomolecular sensor that recognizes ubiquitylated cargo and facilitates their internalization by selectively stabilizing or destabilizing the assembly of endocytic vesicles (Figure 5d-e).

Our study provides new insights into the ubiquitin-dependent role of Epsin1 during CME. Epsin1’s ability to switch from destabilizing Eps15 condensation in ubiquitin’s absence to stabilizing condensation in ubiquitin’s presence reveals a regulatory mechanism that is rooted in the recently discovered ability of endocytic proteins to form multi-valent, condensate-like assemblies in diverse systems including mammalian, yeast, and plant cells^14,19,20^. Epsin1’s ability to sense the presence of ubiquitin within condensates and selectively modulate their stability suggests that condensates could play important roles in endocytic dynamics. More broadly, while the study of protein condensation has focused primarily on biomolecular interactions that promote condensation^69–73^, our results suggest that interactions that destabilize condensation, while little explored, could be equally important to cell physiology and may facilitate environmental sensitivity through switch-like mechanisms.

## Materials and Methods

### Reagents

Tris-HCl (Tris hydrochloride), HEPES (4-(2-hydroxyethyl)-1-piperazineethanesulfonic acid), IPTG (isopropyl-β-D-thiogalactopyranoside), NaCl, β-mercaptoethanol, Triton X-100, neutravidin, and Texas Red-DHPE (Texas Red 1,2-dihexadecanoyl-sn-glycero-3-phosphoethanolamine triethylammonium salt) were obtained from Thermo Fisher Scientific. Sodium bicarbonate, sodium tetraborate, EDTA (Ethylene diamine tetraacetic acid), EGTA (Ethylene glycol tetraacetic acid), glycerol, TCEP (tris(2-carboxyethyl) phosphine), DTT (Dithiothreitol), thrombin, imidazole, sodium bicarbonate, PLL (poly-l-lysine), Atto640 NHS ester, and Atto488 NHS ester were sourced from Sigma-Aldrich. Human Holo-Transferrin protein, Monoubiquitin, and K63 linked Tetraubiquitin (TetraUb) were procured from Boston Biochem (Catalog #: 2914-HT-100MG, U-100H, and UC-310). PEG8K (Polyethylene glycol 8000) was acquired from Promega (Catalog #: V3011). Amine-reactive PEG (mPEG-succinimidyl valerate MW 5000) and PEG-biotin (Biotin-PEG SVA, MW 5000) were purchased from Laysan Bio. DP-EG10-biotin (dipalmitoyl-decaethylene glycol-biotin) was supplied by D. Sasaki (Sandia National Laboratories). POPC (1-palmitoyl-2-oleoyl-glycero-3-phosphocholine) and DGS-NTA-Ni (1,2-dioleoyl-sn-glycero-3-[(N-(5-amino-1-carboxypentyl) iminodiacetic acid)-succinyl] (nickel salt)) were obtained from Avanti Polar Lipids. K63 Tetra-Ubiquitin was sourced from South Bay Bio (SBB-UP0073). siRNA against Epsin1 was sourced from ThermoFisher scientific (HSS121071). All reagents were used without further purification.

### Plasmids and cloning

The pGex4T2 plasmid containing the rat Epsin1 was a gift from H. McMahon^74^. To generate the Epsin1ΔUIM (pGEX4T2-GST-Epsin1ΔUIM) amino acid residues 183-227 were deleted. For Epsin1ΔNPF (pGEX4T2-GST-Epsin1ΔNPF) mutant acid residues 477-551 were deleted. Both of the constructs were custom-designed and constructed by GenScript ([GenScript](https://www.genscript.com/)). The plasmid used for purifying Eps15 was pET28a-6×His-Eps15 encoding H. sapiens Eps15, kindly provided by T. Kirchhausen^75^. Plasmids used for mammalian cell expression of Eps15 variants were derived from Eps15-mCherryN1 (Addgene plasmid #27696, a gift from C. Merrifield^3^, which encodes Eps15-mCherry. All Eps15 variants contain mCherry at their C terminal end for visualization even though mCherry is not mentioned in their names. Plasmids encoding the broad-spectrum deubiquitylase (DUB), UL36, and the catalytically inert mutant (C56S, DUB-dead), were generously provided by J. A. MacGurn^56^. Plasmids encoding the Eps15-DUB and Eps15-DUB-dead were generated by restriction cloning as previously described^15^.

### Protein expression and purification

Epsin1 and its constructs were purified as previously described^76^. In short Epsin1 and its constructs were expressed overnight at 18°C and purified from bacterial extracts by incubation with glutathione–Sepharose beads, followed by extensive washing with 25 mM HEPES, at pH 7.4, 150–300 mM NaCl, 2 mM EDTA and 2 mM dithiothreitol (final buffer composition: 25 mM HEPES at pH 7.4, 150 mM NaCl, 2 mM EDTA and 2 mM dithiothreitol). The protein was cleaved by incubation for 2 h at room temperature and overnight at 4°C with thrombin (Novagen). Thrombin was removed by incubation with benzamidine Sepharose (GE Healthcare) for 30 min at room temperature. Eluted proteins were concentrated and dialysed for 2 h at room temperature in 1L and overnight in 2L of 25 mM HEPES, at pH 7.4, 150 mM NaCl, 1 mM EDTA and 1 mM β-mercaptoethanol.

Eps15 was purified based on a previously reported protocol^14^. Briefly, Eps15 was expressed as N-terminal 6×His-tagged constructs in BL21 (DE3) cells. Protein expression was induced with 1 mM IPTG at 30°C for 6-8 hours. Cells were lysed in a lysis buffer using homogenization and probe sonication on ice. Lysis buffer consisted of 20 mM Tris-HCl, pH 8.0, 300 mM NaCl, 5% glycerol, 10 mM imidazole, 1 mM TCEP, 1 mM PMSF, 0.5% Triton X-100. Eps15 was first purified using Ni-NTA Agarose (Qiagen, Cat#30230) resin. Then it was further purified by gel filtration chromatography using a Superose 6 column run in 20 mM Tris-HCl, pH 8.0, 150 mM NaCl, 1 mM EDTA, and 5 mM DTT. For condensate experiments, prior to running the gel filtration column, the 6×His tag on the proteins was further cleaved with Thrombin CleanCleave kit (Sigma-Aldrich, Cat# RECMT) overnight at 4°C on the rocking table after desalting in 50 mM Tris-HCl pH 8.0, 10 mM CaCl2, 150 mM NaCl and 1 mM EDTA using a Zeba Spin desalting column (Thermo Scientific, Cat#89894).

### Protein labeling

Eps15 was labeled with amine-reactive NHS ester dyes Atto488 in phosphate-buffered saline (PBS, Hyclone) containing 10 mM sodium bicarbonate, pH 8. Epsin1 and its constructs were labeled with amine-reactive NHS ester dyes Atto594 in PBS containing 10 mM sodium bicarbonate, pH 8. TetraUb was labeled with Atto640 in HEPES, pH 7.5. The concentration of dye was adjusted experimentally to obtain a labeling ratio of 0.5–1 dye molecule per protein, typically using 1.5 to 2-fold molar excess of dye. Reactions were performed for 30 min on ice. Then labeled proteins were buffer exchanged into 20 mM Tris-HCl pH 7.5, 150 mM NaCl, 5 mM TCEP, 1 mM EDTA, 1 mM EGTA and separated from unconjugated dye using dialysis (overnight at 4°C). The labeled TetraUb was buffer exchanged to 20 mM Tris-HCl, 150 mM NaCl, pH 7.5 and separated from unconjugated dye as well using 3K Amicon columns. Labeled proteins were dispensed into small aliquots, flash frozen in liquid nitrogen and stored at -80°C. For all condensate experiments a mix of 95% unlabeled and 5% labeled protein was used. 100% labeled TetraUb were directly used due to their small fraction compared to Eps15 variants.

### PLL-PEG preparation

PLL-PEG and biotinylated PLL-PEG were prepared as described previously^15^. Briefly, for PLL-PEG, amine-reactive mPEG-succinimidyl valerate was mixed with poly-L-lysine (15– 30 kD) at a molar ratio of 1:5 PEG to poly-L-lysine. For biotinylated PLL-PEG, amine reactive PEG and PEG-biotin was first mixed at a molar ratio of 98% to 2%, respectively, and then mixed with PLL at 1:5 PEG to PLL molar ratio. The conjugation reaction was performed in 50 mM sodium tetraborate pH 8.5 solution and allowed to react overnight at room temperature with continued stirring. The products were buffer exchanged into 5 mM HEPES, 150 mM NaCl pH 7.4 using Zeba spin desalting columns (7K MWCO, ThermoFisher) and stored at 4°C.

### Protein Condensates

Eps15 condensates were formed by mixing proteins with 3 weight% PEG8K in 20 mM Tris-HCl pH 7.5, 150 mM NaCl, 5 mM TCEP 1 mM EDTA, 1 mM EGTA. 16 μM Eps15 or was used to form the condensates, with addition of Epsin1, and its constructs. For imaging 2% PLL-PEG were used to passivate coverslips (incubated for 20 min) before adding protein-containing solutions. Imaging wells consisted of 5 mm diameter holes in 0.8 mm thick silicone gaskets (Grace Bio-Labs). Gaskets were placed directly onto no.1.5 glass coverslips (VWR International), creating a temporary water-proof seal. Prior to well assembly, gaskets and cover slips were cleaned in 2% v/v Hellmanex III (Hellma Analytics) solution, rinsed thoroughly with water, and dried under a nitrogen stream. The imaging well was washed 6-8 times with 20 mM Tris-HCl, 150 mM NaCl and 5 mM TCEP buffer before adding solutions that contained proteins.

### GUV preparation

GUVs consisted of 80 mol% POPC, 8 mol% DOPS, 5 mol% DGS-NTA-Ni, 5 mol% PIP2 (L-α-phosphatidylinositol-4,5-bisphosphate) and 2 mol% DP-EG10-biotin. GUVs were prepared by electroformation according to published protocols^14^. Briefly, lipid mixtures dissolved in chloroform were spread into a film on indium-tin-oxide (ITO) coated glass slides (resistance ∼8-12 W per square) and further dried in a vacuum desiccator for at least 2 hours to remove all of the solvent. Electroformation was performed at 55°C in glucose solution with an osmolarity that matched the buffer to be used in the experiments. The voltage was increased every 3 min from 50 to 1400 mV peak to peak for the first 30 min at a frequency of 10 Hz. The voltage was then held at 1400 mV peak to peak, 10 Hz, for 120 min and finally was increased to 2200 mV peak to peak, for the last 30 min during which the frequency was adjusted to 5 Hz. GUVs were used on the same day after electroformation.

### GUV tethering

GUVs were tethered to glass coverslips for imaging as previously described^77^. Briefly, glass cover slips were passivated with a layer of biotinylated PLL-PEG, using 5 kDa PEG chains. GUVs doped with 2 mol% DP-EG10-biotin were then tethered to the passivated surface using neutravidin. In each imaging well, 20 μL of biotinylated PLL-PEG was added. After 20 min of incubation, wells were serially rinsed with appropriate buffers by gently pipetting. Next, 4 μg of neutravidin dissolved in 25 mM HEPES, 150 mM NaCl (pH 7.4) was added to each sample well and allowed to incubate for 10 minutes. Wells were then rinsed with the appropriate buffer to remove excess neutravidin. GUVs were diluted appropriately in 20 mM Tris-HCl, 150 mM NaCl, 5 mM TCEP, pH 7.5 and allowed to incubate for 10 minutes. Excess GUVs were then rinsed from the well using the same buffer and the sample was subsequently imaged using confocal fluorescence microscopy.

### Cell culture

Human-derived SUM159 cells gene-edited to add a HaloTag to both alleles of AP2 σ2 were a gift from T. Kirchhausen^50^. Cells were further gene-edited to knock out both alleles of endogenous Eps15 using CRISPR-associated protein 9 (Cas9) to produce the Eps15 knock out cells developed previously by our group^14^. Cells were grown in 1:1 DMEM high glucose: Ham’s F-12 (Hyclone, GE Healthcare) supplemented with 5% fetal bovine serum (Hyclone), Penicillin/Streptomycin/l-glutamine (Hyclone), 1 μg ml^−1^ hydrocortisone (H4001; Sigma-Aldrich), 5 μg ml^−1^ insulin (I6634; Sigma-Aldrich) and 10 mM HEPES, pH 7.4 and incubated at 37°C with 5% CO_2_. Cells were seeded onto acid-washed coverslips. Plasmid transfection was carried out by mixing 1μg of plasmid DNA with 3 μl Fugene HD transfection reagent (Promega). HaloTagged AP2 σ2 was visualized by adding Janelia Fluor 646-HaloTag ligand (Promega).

### siRNA transfection and Epsin1 knock down

Approximately 60,000 cells were seeded per well in a 6-well plate and incubated for 24 hours prior to transfection. siRNA knock down was performed using Lipofectamine RNAiMAX (ThermoFisher Scientific, catalog no. 13778075). For the transfection mixture, 6.5 µl of Lipofectamine RNAiMAX and 5.5 µl of 20 µM siRNA were each diluted in 100 µl of OptiMEM and incubated separately at room temperature for 5 minutes. The siRNA solution was then combined with the Lipofectamine RNAiMAX solution and incubated for an additional 8–10 minutes at room temperature before being added dropwise to the cells in fresh medium. Transfections were conducted twice, with a 24-hour interval between rounds, and measurements were taken on the fourth day after cell plating. Western blot analysis confirmed knock down efficiencies exceeding 82% for all targeted proteins. Control cells were transfected concurrently with control siRNA obtained from QIAGEN (Germantown, MD).

### Western blot analysis

Knock down of Epsin1 was verified through Western blot analysis. siRNA-treated SUM cells were washed with ice-cold Dulbecco’s Phosphate Buffered Saline (Sigma) and lysed in 1% NP-40 lysis buffer (100 mM Tris, pH 7.5, 100 mM NaCl, 1% NP-40) supplemented with 170 mg/mL PMSF, 2 mg/mL leupeptin, 2 mg/mL aprotinin, and 10 mM DTT. Protein concentration in the lysates was determined using the Coomassie Protein Assay Reagent (ThermoScientific), and 30 µg of protein per sample was standardized for loading. The lysates were mixed with 4x SDS-PAGE loading buffer (250 mM Tris, pH 6.8, 400 mM DTT, 40% glycerol, 8% SDS, 60 µM bromophenol blue), boiled at 100°C for 5 minutes, and resolved on a 4–12% BOLT Bis-Tris Plus polyacrylamide gel (Invitrogen). Proteins were transferred to a nitrocellulose membrane (BioRad), blocked for 1 hour at room temperature with LI-COR Block Buffer, and probed with an anti-Epsin1 antibody (mouse polyclonal, Santa Cruz) diluted 1:1000 in 1x TNET Wash Buffer (10 mM Tris, pH 7.5, 2.5 mM EDTA, pH 8.0, 50 mM NaCl, 0.1% Tween-20). After three washes in TNET Wash Buffer (5 minutes each), the membrane was incubated for 1 hour at room temperature with secondary Goat anti-Mouse IgG (H+L) Alexa Fluor™ Plus 680 antibody (Invitrogen) diluted 1:15,000 in Intercept Antibody Diluent Buffer (LI-COR). Following another three washes in TNET Wash Buffer, the blot was imaged using a LI-COR Odyssey Imaging System. The membrane was subsequently reprobed with anti-β Actin antibody and the same secondary antibody before imaging again.

### Fluorescence microscopy

Images of protein condensates and GUVs were collected on a spinning disc confocal super resolution microscope (SpinSR10, Olympus, USA) equipped with a 1.49 NA/100X oil immersion objective. For GUV imaging, image stacks taken at fixed distances perpendicular to the membrane plane (0.5 μm steps) were acquired immediately after GUV tethering and again after protein addition. Images taken under deconvolution mode were processed by the built-in deconvolution function in Olympus CellSens software (Dimension 3.2, Build 23706). For condensates imaging was performed 5 min after adding proteins, providing sufficient time to achieve protein binding and reach a steady state level of binding. For experiments used to construct phase diagrams, temperature was monitored by a thermistor placed in the protein solution in a sealed chamber to prevent evaporation. Samples were heated from room temperature through an aluminum plate fixed to the top of the chamber. Temperature was increased in steps of 1°C (and was held onto the temperature for 1 minute) until the critical temperature was reached. Live cell plasma membrane imaging was conducted using the same Olympus spinning disk confocal microscope. Fluorescence emission was captured using a Hamamatsu ORCA C13440-20CU CMOS camera. Excitation was achieved with lasers at 488 nm for GFP, and 640 nm for JF646.

Live-cell plasma membrane images were collected on a TIRF microscope consisting of an Olympus IX73 microscope body, a Photometrics Evolve Delta EMCCD camera, and an Olympus 1.4 NA ×100 Plan-Apo oil objective, using MicroManager version 1.4.23. All live-cell imaging was conducted in TIRF mode at the plasma membrane 24 - 48 hours after transfection. Imaging media was supplemented with 1 μL OxyFluor (Oxyrase, Mansfield, OH) per 33 μL media to decrease photobleaching during live-cell fluorescence imaging. The coverslip was heated to produce a sample temperature of 37°C using an aluminum plate fixed to the back of the sample. 532 nm and 640 nm lasers were used for excitation of mCherry and JF646-HaloTag ligands of AP2, respectively. Cell movies were collected over 10 min at 2 s intervals between frames.

### Flow cytometry

The internalization of transferrin (Tf) through clathrin-mediated endocytosis was measured using flow cytometry based on the previous report^59^. First, holo-transferrin was labeled with Atto-488. SUM cells were plated on a 6-well plate at a density of 100,000 cells per well and a total volume of 2 mL per well. After 24 hr, media was aspirated from the cells and they were rinsed with PBS 3 times. The cells were starved for 30 min in 1mL pre-warmed serum-free media and were then incubated with 50 μg/mL Tf-Atto 488 for 25 min. Subsequently, the cells were incubated with 500 μL trypsin for about 2 min at 37°C, and then the trypsin was removed and replaced with fresh trypsin. This process was repeated twice to wash away excess Tf. Then cells were fully detached from the wells by incubation with trypsin for 5 min. The trypsin was then quenched with 1 mL of media, and cells from each well were collected and resuspended in 300 μL of cold PBS in preparation for flow cytometry analysis.

A Guava easyCyte Flow Cytometer (Millipore Sigma) with 488 nm and 532 nm excitation lasers was used to analyze fluorescence of cells after Tf-Atto 488 incubation. All data were collected at 35 μL/min and flow cytometry data were analyzed using FlowJo (Treestar). A gate was drawn in forward scattering versus side scattering plots to exclude debris. Within this gate, cell populations were further analyzed by plotting histograms of FITC fluorescence intensity to determine shifts in Tf-Atto 488 uptake for each group. Mean fluorescence intensity was used to indicate the level of transferrin internalization.

### Image analysis

Fluorescence images were analyzed in ImageJ (http://rsbweb.nih.gov/ij/). Intensity values along line scans were measured in unprocessed images using ImageJ. For phase diagrams, fluorescence intensity was measured in the center square of a 3 × 3 grid for each image where illumination was even. The background intensity was subtracted in all images. Clathrin-coated structures were detected and tracked using CME analysis in MATLAB^9^. The point spread function of the data was used to determine the standard deviation of the Gaussian function. AP2 σ2 signal was used as the master channel to track clathrin-coated structures. Detected structures were analyzed if they persisted in at least three consecutive frames. The lifetimes of clathrin-coated pits that met these criteria were recorded for lifetime distribution analysis under different conditions.

### Statistical analysis

For all experiments yielding micrographs, each experiment was repeated independently on different days at least three times (n = 3), with similar conditions. Collection of cell image data for CME analysis was performed independently on at least two different days for each cell type and n= 15 to 20 independent cells were used for each group. More than 10000 pits were analyzed for each condition. Statistical analysis was carried out using a two-tailed student’s t-test (unpaired, unequal variance) on GraphPad Prism to probe for statistical significance (p < 0.05).

## Acknowledgements

We thank T. Kirchhausen (Harvard U) for the gift of SUM159/AP2σ2-HaloTag cells and J. MacGurn (Vanderbilt U) for the gift of plasmids encoding deubiquitylase UL36 and its catalytically inactive mutant. We acknowledge funding from NIH/NIGMS (R35GM139531) and NSF BIO (1934509) to J. C. Stachowiak.

## Contributions

S.S., E.M.L., J.M.H., and J.C.S. designed experiments. L.W. and E.M.L. purified Eps15, Epsin1 and the variants for *in vitro* experiments; S.S. J.M.H., and J.C.S. wrote and edited the manuscript. S.S., HY.L., J.P., F.Y., L.W., E.M.L., J. M. H., and J.C.S. performed experiments and analysed data. All authors consulted on manuscript preparation and editing.

## Supplementary Material

This PDF file includes Supplementary Figures S1 to S13 and Supplementary Movie Captions S1 to S3.

**Figure S1.**
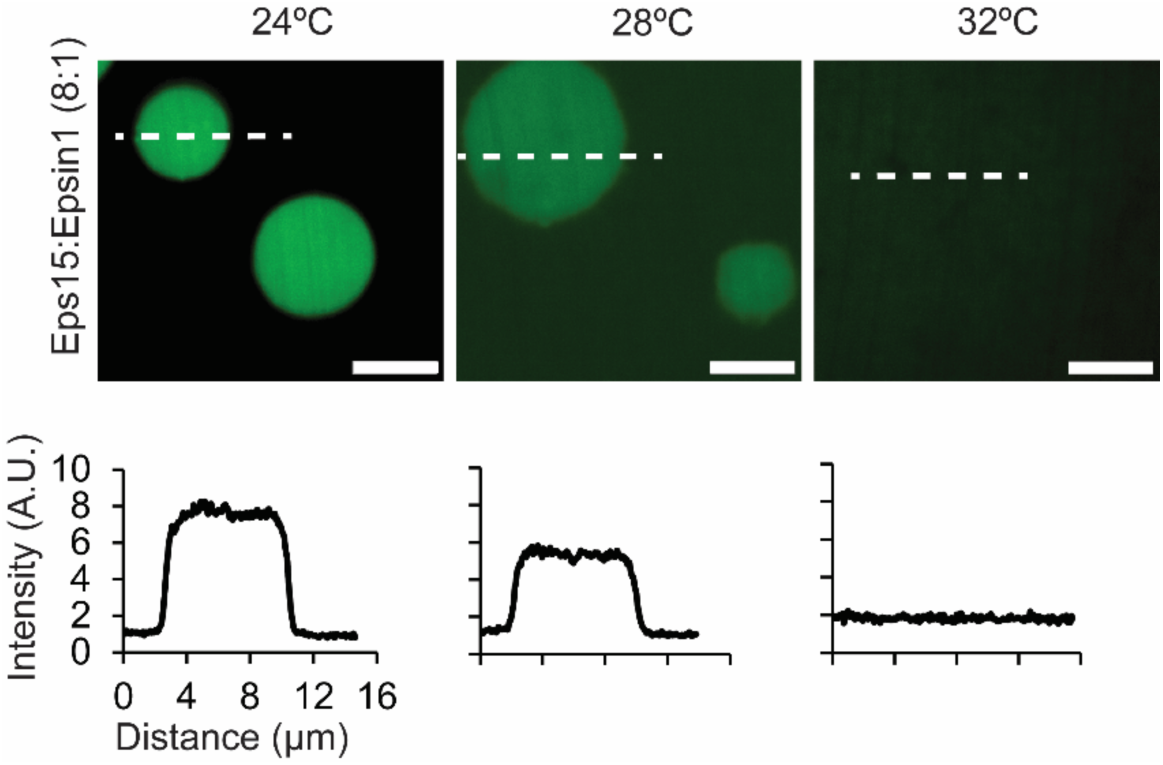
Representative images of protein condensates composed of Eps15 to Epsin1 in 8:1 ratio at increasing temperatures. Plots show fluorescence intensity of Eps15 measured along dotted lines in each image. Condensates are formed from 16 μM Eps15 mixed with 2 μM Epsin1 in 20 mM Tris-HCl, 150 mM NaCl, 5 mM TCEP, 1 mM EDTA and 1 mM EGTA at pH 7.5 buffer with 3% PEG8K. n = 3 biological replicates, scale bars equal 5 μm.

**Figure S2.**
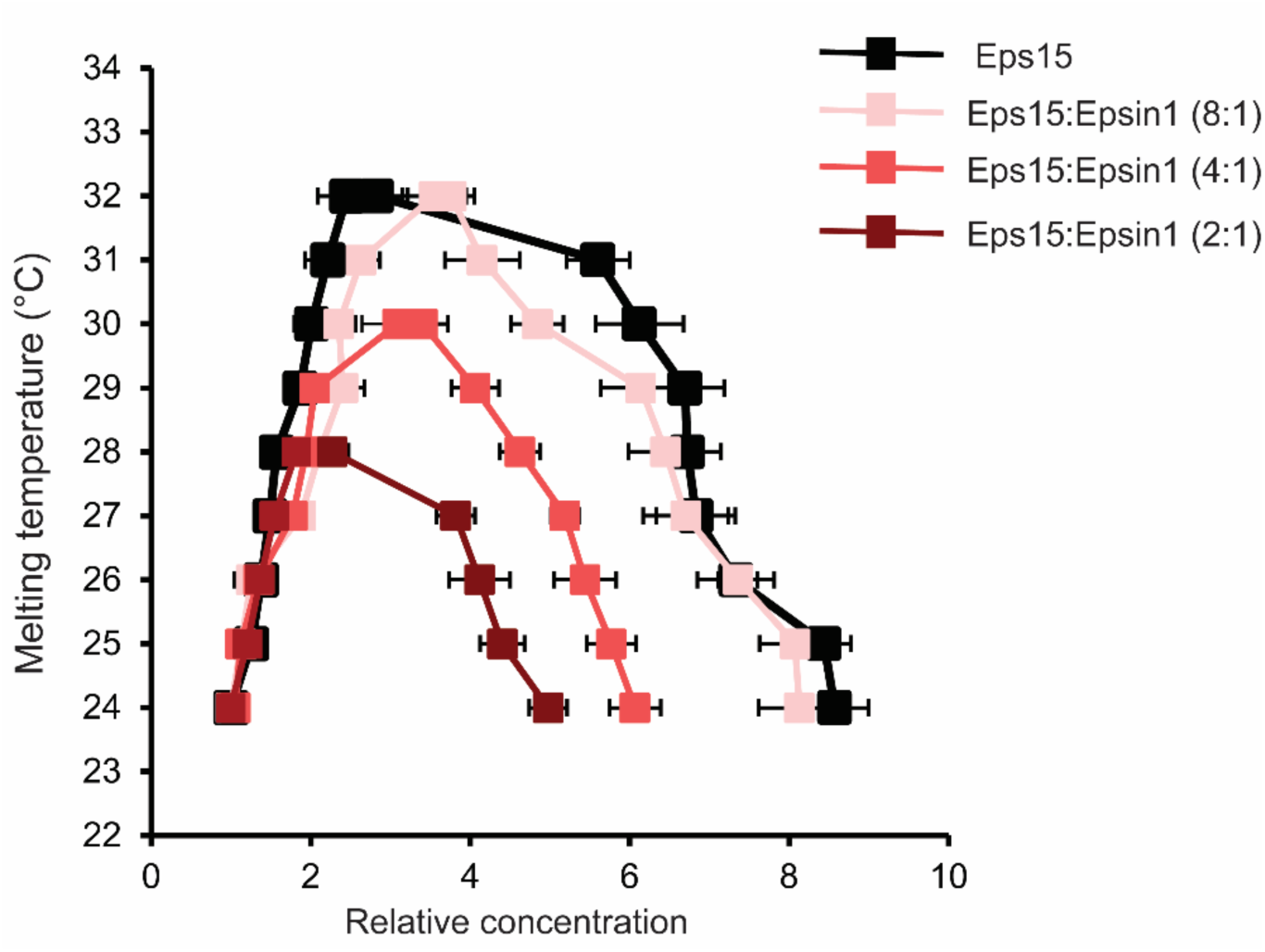
Phase diagram of Eps15 condensates and Eps15:Epsin1 mixed condensates at 8:1, 4:1 and 2:1 ratio. Condensates mapped by Atto488-labelled Eps15 fluorescence intensity. Intensity was normalized based on the intensity of the solution. Dots on the right side are protein concentrations in condensates and dots on the left side are concentrations in solution. At least 20 condensates are analyzed under each temperature. Data are mean ± SD. Condensates are formed in 20 mM Tris-HCl, 150 mM NaCl, 5 mM TCEP, 1 mM EDTA and 1 mM EGTA at pH 7.5 buffer with 3% PEG8K. n = 3 biological replicates.

**Figure S3.**
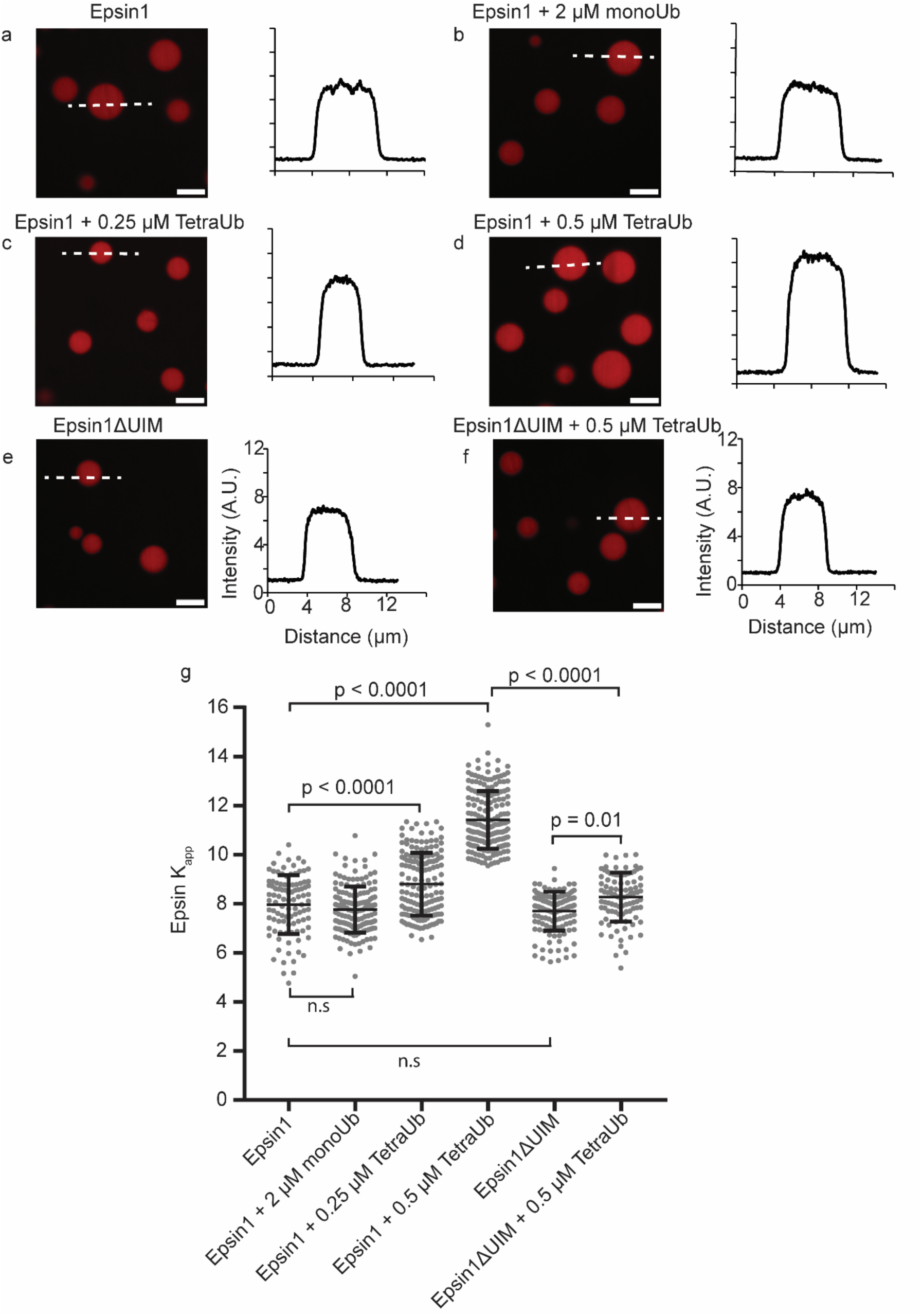
TetraUb mediated cross-linking between Eps15 and Epsin1. **a-f,** Representative images showing the Epsin1 and Epsin1ΔUIM partition (red) in Eps15 condensates incubated with and without TetraUb and monoUb. Plots on the right depict intensity profiles of Epsin1 and Epsin1ΔUIM along the white dashed line shown in the corresponding images. Epsin1 partition in Eps15 condensates **(a)**. Epsin1 partition in Eps15 condensates in presence of 2 μM monoUb **(b)**. Epsin1 partition in Eps15 condensates in presence of 0.25 μM TetraUb **(c)**. Epsin1 partition in Eps15 condensates in presence of 0.5 μM TetraUb **(d)**. Epsin1ΔUIM partition in Eps15 condensates **(e)**. Epsin1ΔUIM partition in Eps15 condensates in presence of 0.5 μM TetraUb **(f)**. **g**, Plot shows Epsin1/Epsin1ΔUIM partition coefficient (Kapp) in Eps15 condensates in presence of monoTetraUb. All condensate experiments were performed by mixing 1.6 μM Epsin1 or Epsin1ΔUIM with 16 μM Eps15 (unlabeled) (1:10 ratio) in 20 mM Tris-HCl, 150 mM NaCl, 5 mM TCEP, 1 mM EDTA and 1 mM EGTA at pH 7.5 with 3% w/v PEG8K. The increased Kapp in Eps15 condensates in presence of TetraUb was found to be dependent on the UIM region of Epsin1. n = 3 biologically independent experiments with at least 100 total condensates were measured for each condition. Data are mean ± SD. An unpaired, two tailed student’s t test was used for statistical significance using GraphPad Prism. Imaging was carried out at room temperature, 24°C. All scale bars equal 5 μm.

**Figure S4.**
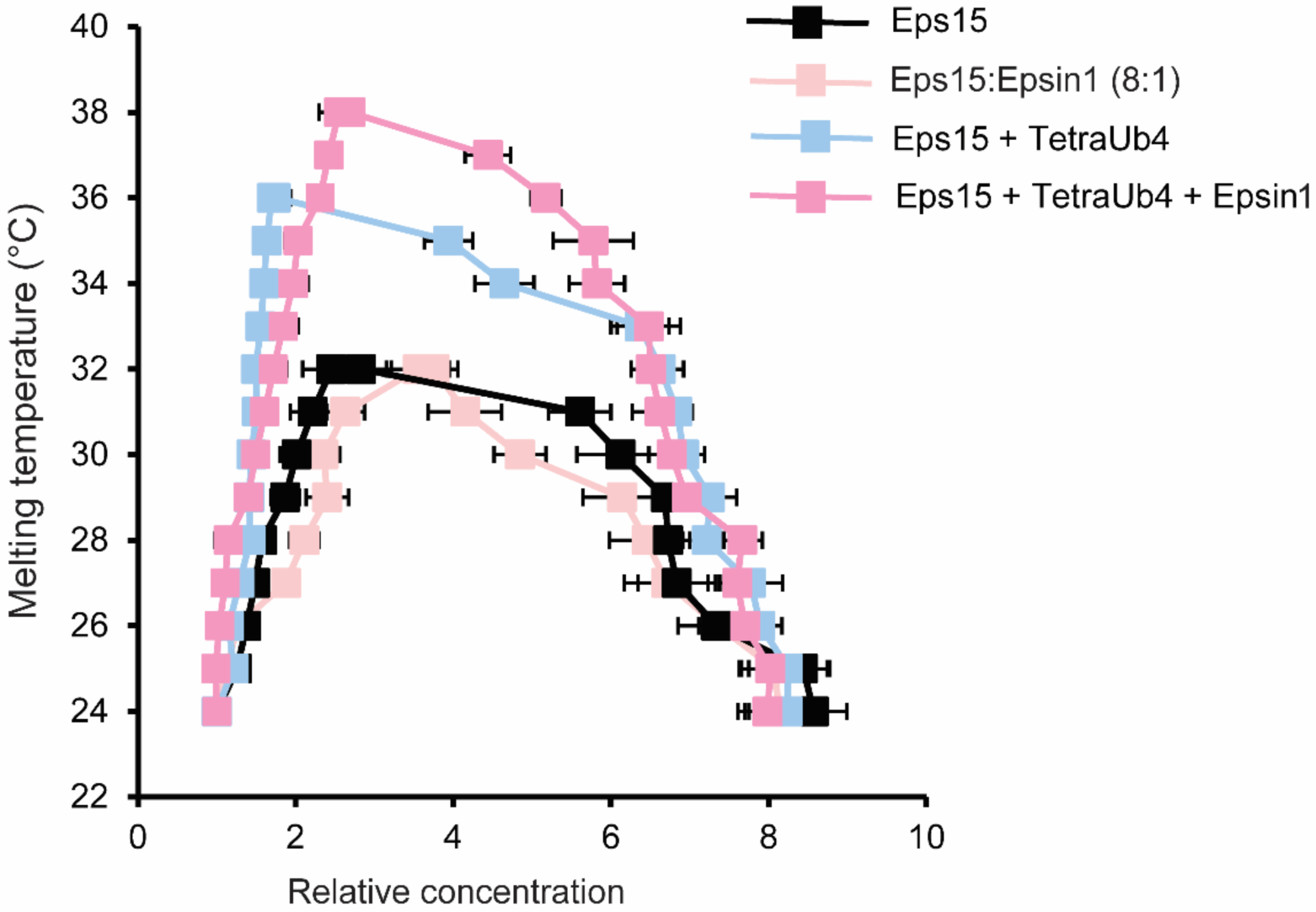
Phase diagram (Tm) of condensates consisting of Eps15 alone, Eps15:Epsin1 mixed at 8:1 ratio, Eps15 mixed with 0.12 μM TetraUb and Eps15:Epsin1 8:1 system incubated with 0.12 μM TetraUb. Condensates mapped by Atto488-labelled Eps15 fluorescence intensity. Intensity was normalized based on the intensity of the solution. Dots on the right side are protein concentrations in condensates and dots on the left side are concentrations in solution. Eps15 alone and Eps15:Epsin1 8:1 both melts at 32°C. Addition of TetraUb in Eps15 condensates increased the Tm to 36°C. Addition of Epsin1 in this system (Eps15 + TetraUb + Epsin1) increased the Tm further to 38°C. At least 20 condensates are analyzed under each temperature. Data are mean ± SD. Condensates are formed in 20 mM Tris-HCl, 150 mM NaCl, 5 mM TCEP, 1 mM EDTA and 1 mM EGTA at pH 7.5 buffer with 3% PEG8K. n = 3 biological replicates.

**Figure S5.**
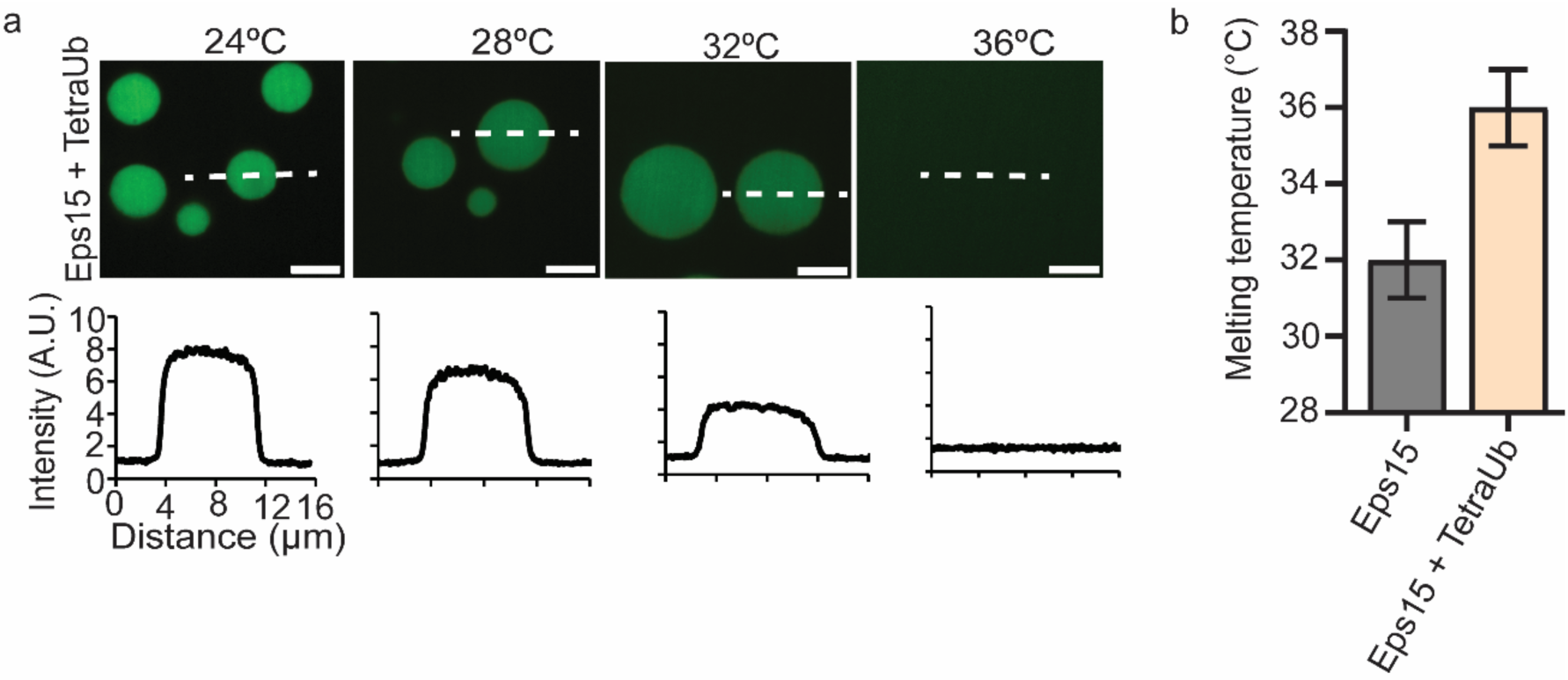
a, Representative images of protein condensates composed of Eps15 and TetraUb at increasing temperatures. The intensity profile below shows fluorescence intensity of Eps15 (Atto 488) measured along dotted lines in each image. **b,** Bar graph shows the melting temperature (Tm) of Eps15 and Eps15-TetraUb mixed condensates, n = 3. The error bar represents ± 1°C of the melting temperature. Condensates are formed in 20 mM Tris-HCl, 150 mM NaCl, 5 mM TCEP, 1 mM EDTA and 1 mM EGTA at pH 7.5 buffer with 3% PEG8K. The scale bars are 5 µm.

**Figure S6.**
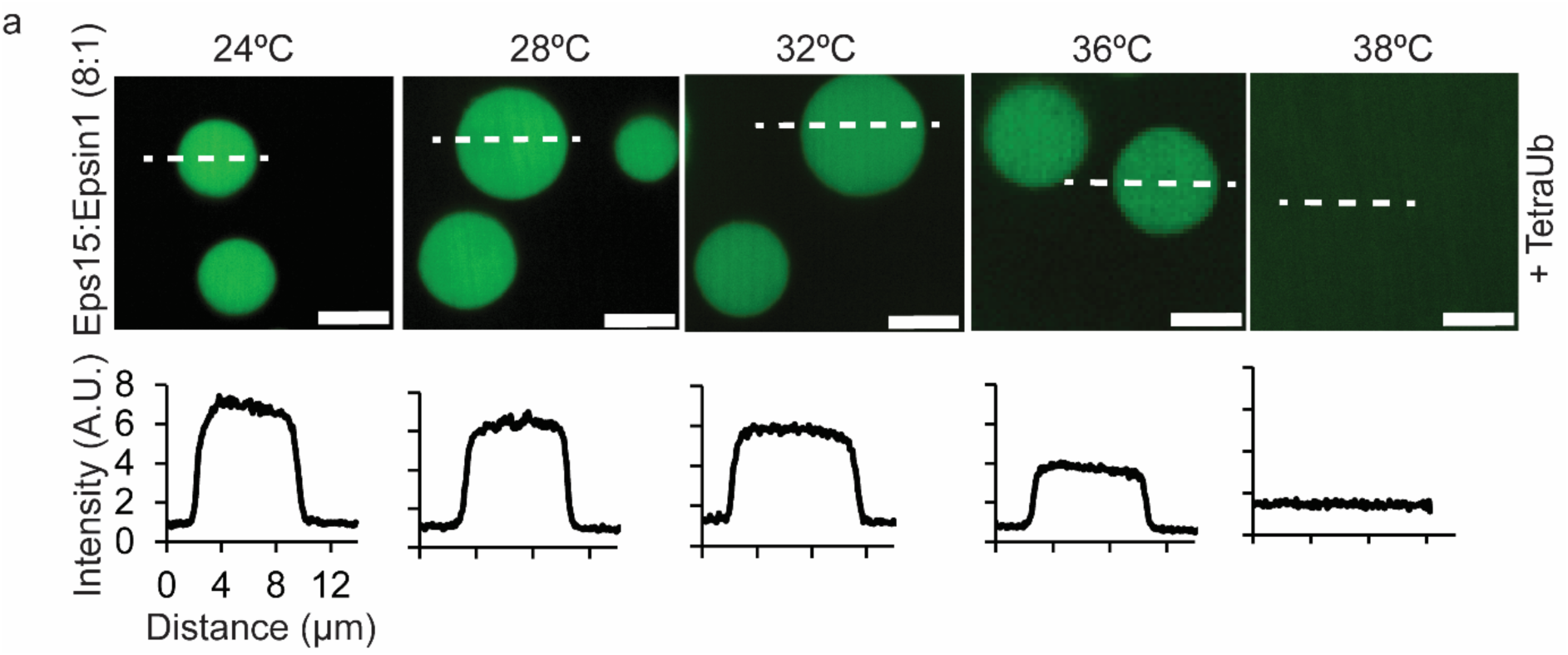
a, Representative images of protein condensates composed of 16 μM Eps15 mixed with 2 μM Epsin1 and 0.12 μM TetraUb at increasing temperatures. Intensity plots show fluorescence intensity of Eps15 measured along dotted lines in each image, n = 3 replicates. All condensate experiments were performed in 20 mM Tris-HCl, 150 mM NaCl, 5 mM TCEP, 1 mM EDTA and 1 mM EGTA at pH 7.5 with PEG8K. All scale bars equal 5 μm.

**Figure S7.**
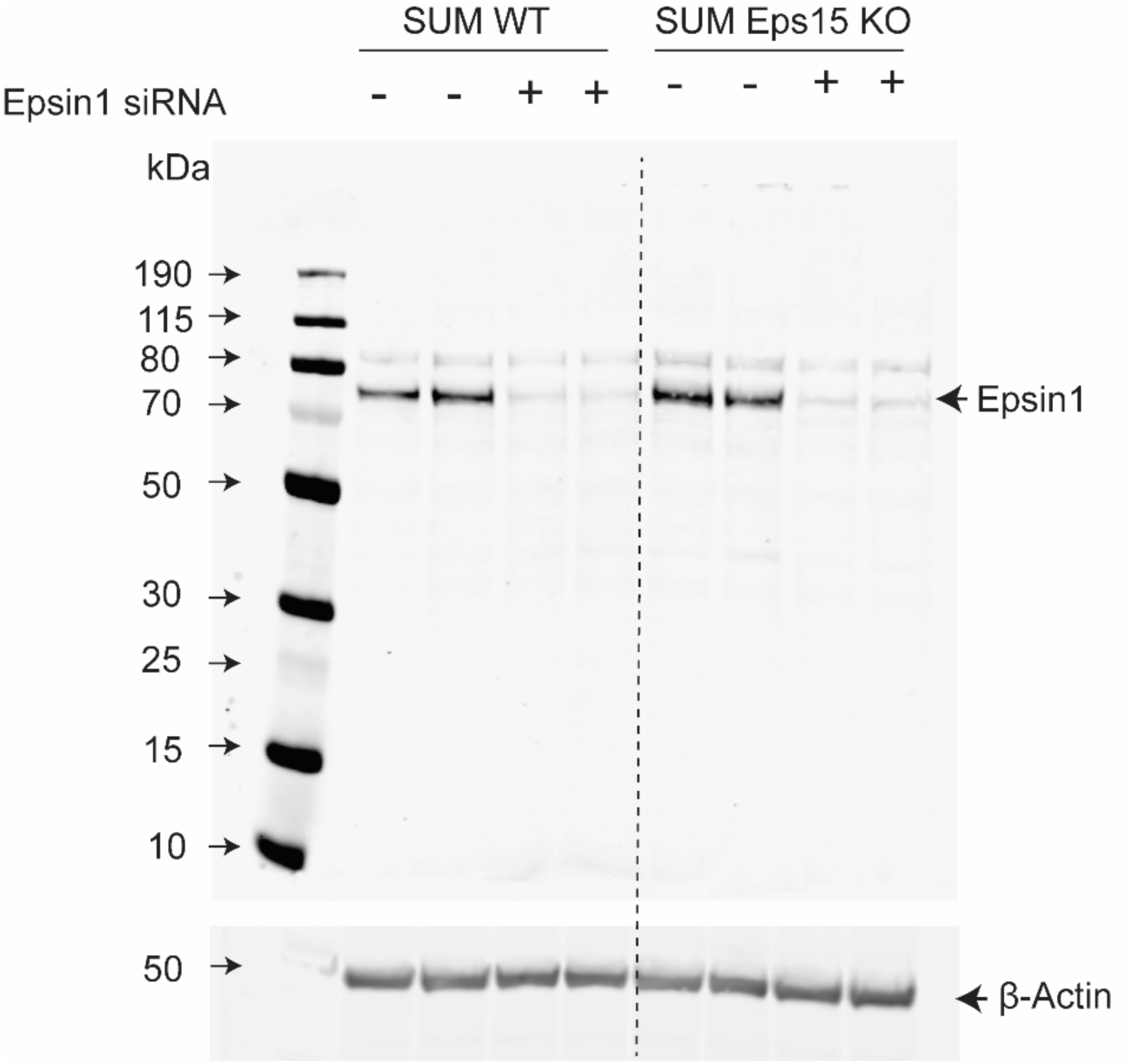
Whole cell lysates from wild type SUM159/AP2-HaloTag (SUM WT) cells and SUM159/AP2-HaloTag Eps15 CRISPR knock out (SUM Eps15 KO) cells were separated by SDS-PAGE and immunoblotted for Epsin1 and *β*-Actin. Both cell types were transfected with siRNA against Epsin1 (twice, 24 hour interval) and were collected 24 hours post 2nd transfection.

**Figure S8.**
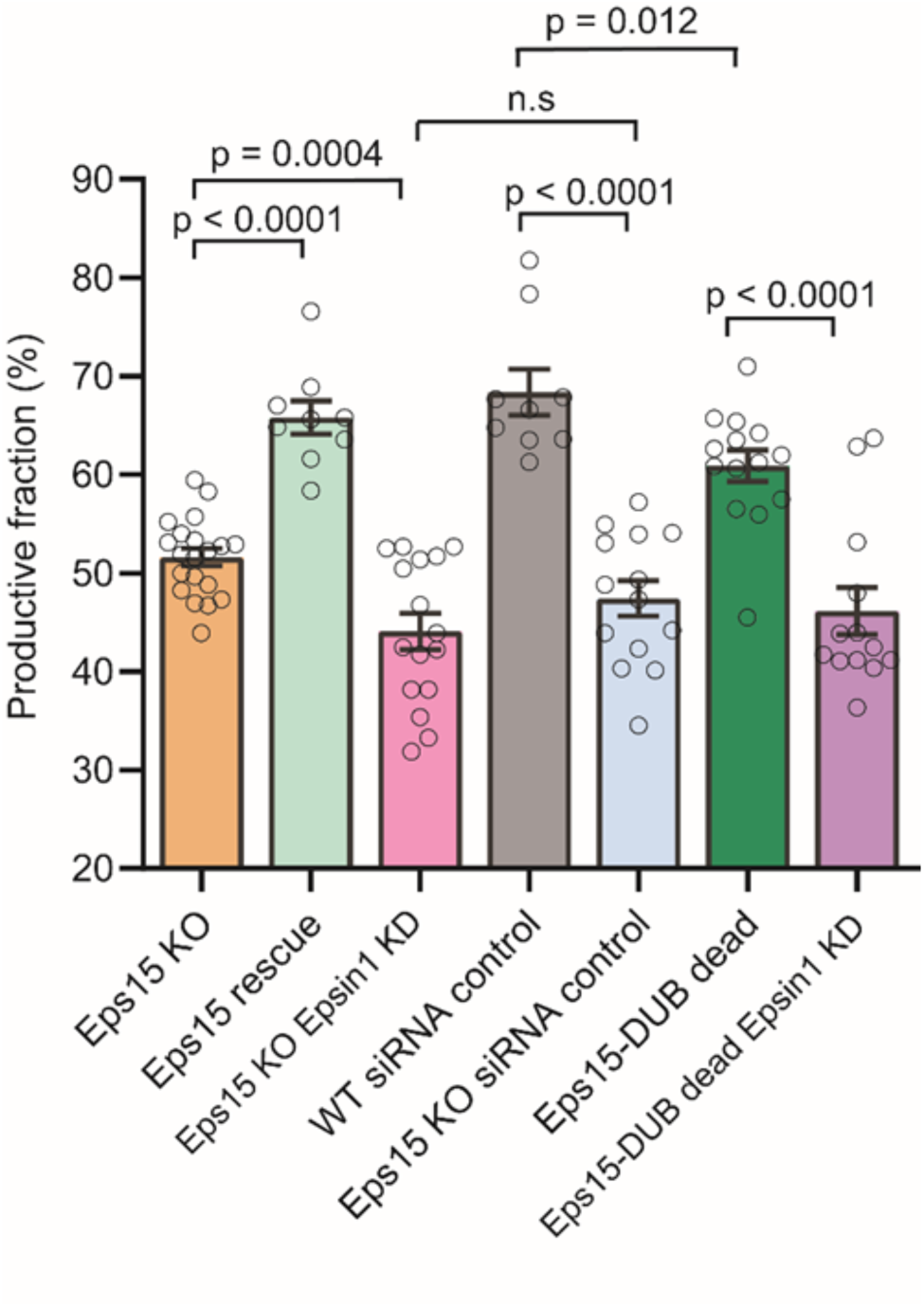
Bar plot of the productive fraction (lifetime between 20 - 180 sec) for all four groups. Data are mean ± S.E.M. Eps15 KO represents SUM cells that were CRISPR modified to knock out alleles of endogenous Eps15. Eps15 rescue represents Eps15 KO cells expressing wild type Eps15. Eps15 KO Epsin1 KD represents Eps15 KO cells also knocked down for Epsin1 using siRNA. WT siRNA control represents wild type cells transfected with control siRNA. Eps15 KO siRNA control represents Eps15 KO cells transfected with control siRNA. Eps15-DUB dead represent Eps15 KO cells transfected with wild type Eps15 fused with catalytically inactive DUB (fused to C terminal end of Eps15). Eps15-DUB dead Epsin1 KD represent Eps15 KO cells transfected with wild type Eps15 fused with catalytically inactive DUB (fused to C terminal end of Eps15) and knocked down for Epsin1 using siRNA. All Eps15 constructs have mCherry at their C terminus for visualization. For Eps15 KO, Eps15 rescue, Eps15 KO Epsin1 KD, WT siRNA, Eps15 KO siRNA, Eps15-DUB dead and Eps15-DUB dead Epsin1 KD n = 19, 10, 16, 10, 14, 15 and 14 biologically independent cells, respectively, were used to collect data. In total > 10000 pits were analyzed for each group. An unpaired, two-tailed student’s t test was used for statistical significance using GraphPad prism, n.s. means no significant difference. p < 0.05 is considered significantly different. All cell images were collected at 37°C.

**Figure S9.**
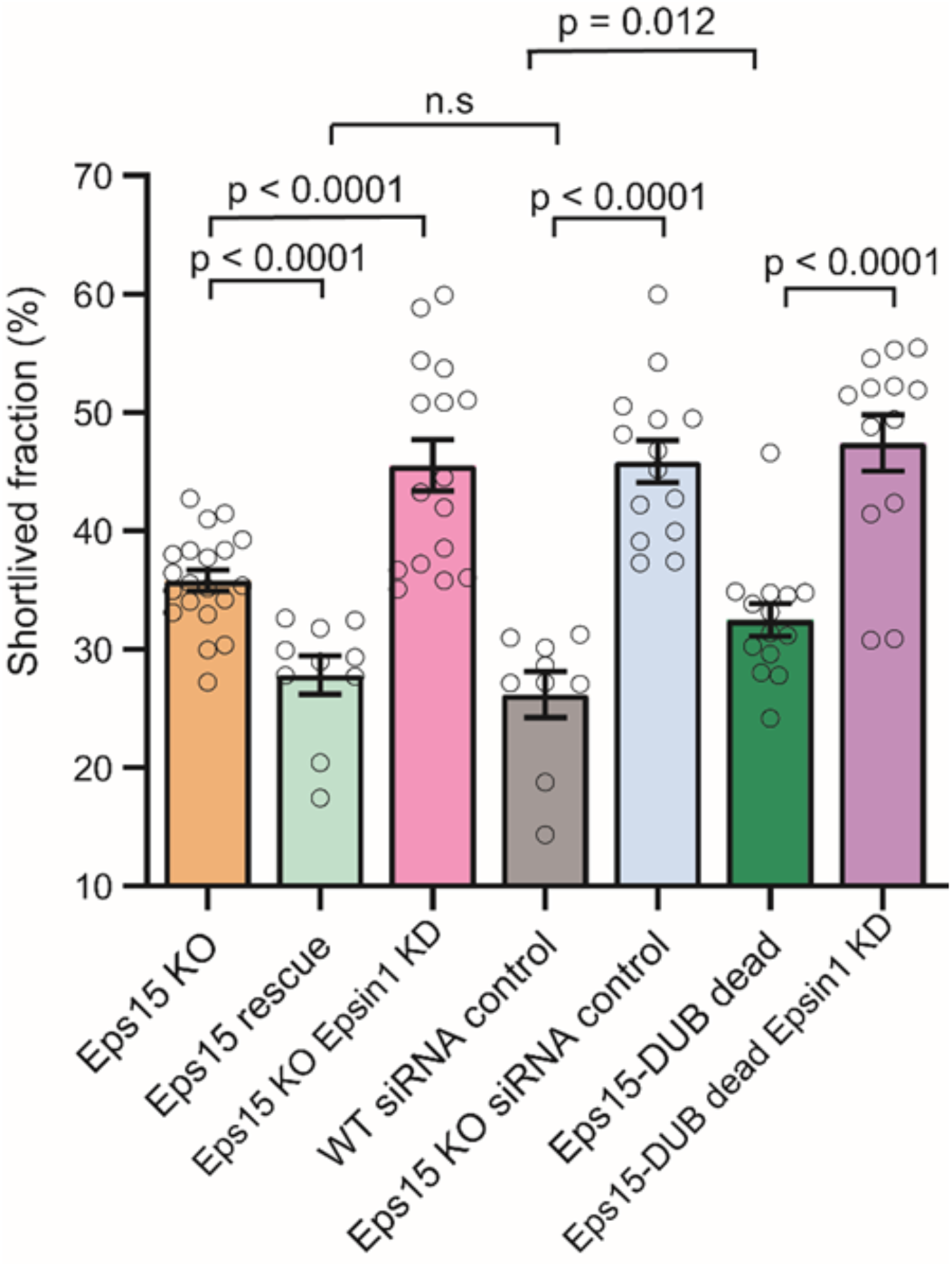
Bar plot of the short-lived fraction (lifetime between < 20 sec) for all four groups. Data are mean ± S.E.M. Eps15 KO represents SUM cells that were CRISPR modified to knock out alleles of endogenous Eps15. Eps15 rescue represents Eps15 KO cells expressing wild type Eps15. Eps15 KO Epsin1 KD represents Eps15 KO cells also knocked down for Epsin1 using siRNA. WT siRNA control represents wild type cells transfected with control siRNA. Eps15 KO siRNA control represents Eps15 KO cells transfected with control siRNA. Eps15-DUB dead represent Eps15 KO cells transfected with wild type Eps15 fused with catalytically inactive DUB (fused to C terminal end of Eps15). Eps15-DUB dead Epsin1 KD represent Eps15 KO cells transfected with wild type Eps15 fused with catalytically inactive DUB (fused to C terminal end of Eps15) and knocked down for Epsin1 using siRNA. All Eps15 constructs have mCherry at their C terminus for visualization. For Eps15 KO, Eps15 rescue, Eps15 KO Epsin1 KD, WT siRNA, Eps15 KO siRNA, Eps15-DUB dead and Eps15-DUB dead Epsin1 KD n = 19, 10, 16, 10, 14, 15 and 14 biologically independent cells, respectively, were used to collect data. In total > 10000 pits were analyzed for each group. An unpaired, two-tailed student’s t test was used for statistical significance using GraphPad prism, n.s. means no significant difference. p < 0.05 is considered significantly different. All cell images were collected at 37°C.

**Figure S10.**
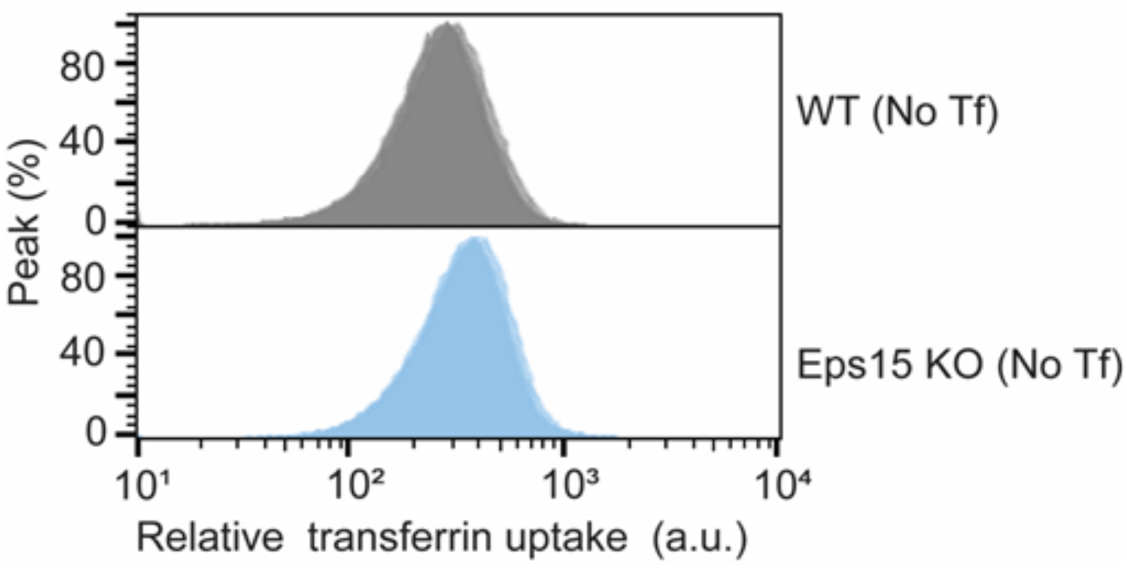
Flow cytometry histogram of the cells in absence of transferrin (Tf), n = 3 biological replicates. Wild type (WT) represents SUM cells that have endogenous Eps15 and Epsin1 expression. Eps15 KO represents SUM cells that were CRISPR modified to knock out alleles of endogenous Eps15. All flow cytometry runs were carried out at 24°C. The measured fluorescence reflects autofluorescence of the cells and background noise levels in the flow cytometry system. These values represent a small contribution (1-2%) of the values for cells exposed to fluorescent Tf (Figure 5 and Figure S11).

**Figure S11.**
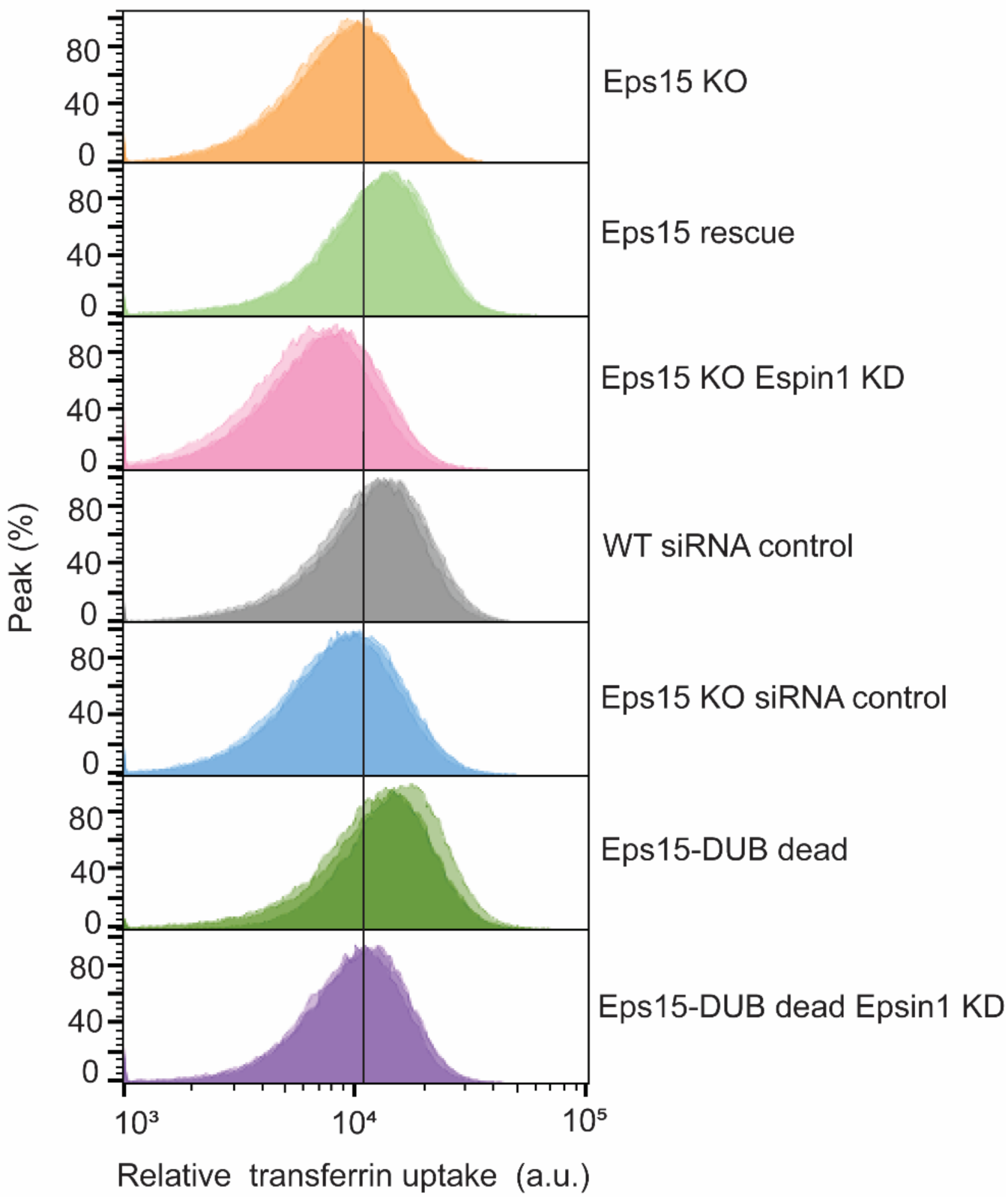
Flow cytometry histogram of the transferrin fluorescence intensity (green fluorescence) of cells in each condition, n = 3 biological replicates. Eps15 KO represents SUM cells that were CRISPR modified to knock out alleles of endogenous Eps15. Eps15 rescue represents Eps15 KO cells expressing wild type Eps15. Eps15 KO Epsin1 KD represents Eps15 KO cells also knocked down for Epsin1 using siRNA. WT siRNA control represents wild type cells transfected with control siRNA. Eps15 KO siRNA control represents Eps15 KO cells transfected with control siRNA. Eps15-DUB dead represent Eps15 KO cells transfected with wild type Eps15 fused with catalytically inactive DUB (fused to C terminal end of Eps15). Eps15-DUB dead Epsin1 KD represent Eps15 KO cells transfected with wild type Eps15 fused with catalytically inactive DUB (fused to C terminal end of Eps15) and knocked down for Epsin1 using siRNA. All Eps15 constructs have mCherry at their C terminus for visualization. All flow cytometry runs were carried out at 24°C.

**Figure S12.**
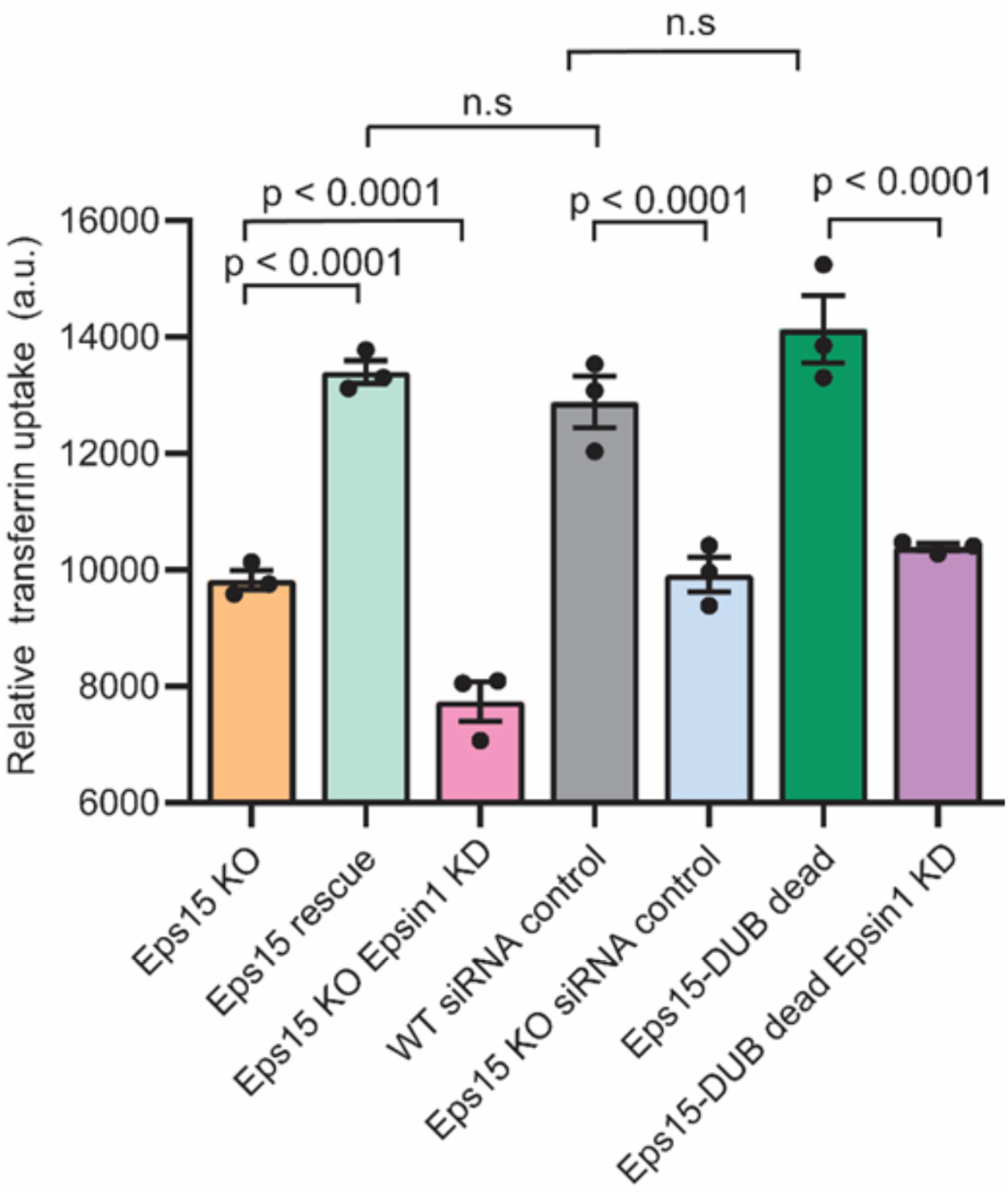
Bar graph represents transferrin fluorescence intensity measured by flow cytometry for all groups. Data are mean ± standard deviation, n = 3 biological replicates. Eps15 KO represents SUM cells that were CRISPR modified to knock out alleles of endogenous Eps15. Eps15 rescue represents Eps15 KO cells expressing wild type Eps15. Eps15 KO Epsin1 KD represents Eps15 KO cells also knocked down for Epsin1 using siRNA. WT siRNA control represents wild type cells transfected with control siRNA. Eps15 KO siRNA control represents Eps15 KO cells transfected with control siRNA. Eps15-DUB dead represent Eps15 KO cells transfected with Eps15 fused with catalytically inactive DUB (fused to C terminal end of Eps15). Eps15-DUB dead Epsin1 KD represent Eps15 KO cells transfected with wild type Eps15 fused with catalytically inactive DUB (fused to C terminal end of Eps15) and knocked down for Epsin1 using siRNA. All Eps15 constructs have mCherry at their C terminus for visualization. flow cytometry runs were carried out at 24°C.

**Figure S13.**
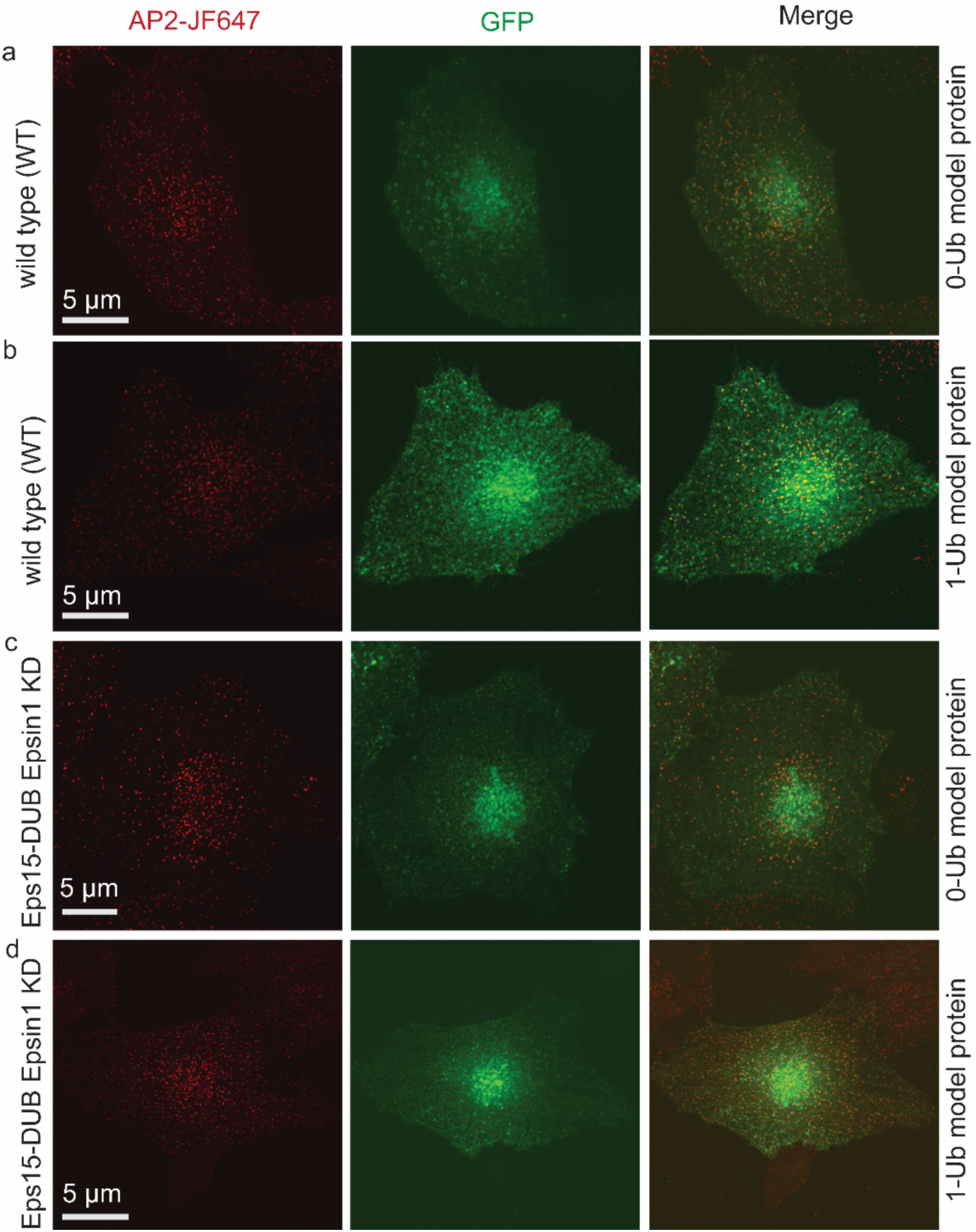
Whole cell fluorescence images of the plasma membrane of SUM cells expressing the model proteins. **a-b**, Spinning disk confocal images of wild type (WT) whole cells expressing the 0-Ub model protein **(a)** and 1-Ub model protein **(b)**. **c-d**, Spinning disk confocal images of Eps15 KO Epsin1 KD whole cells expressing the 0-Ub model protein **(c)** and 1-Ub model protein **(d)**. Crops of these cells are shown in Figure 6 c-d, respectively. Red fluorescence (AP2-JF646) marks endocytic sites and green fluorescence (GFP) highlights the model proteins. The scale bars are 5 µm.

**Movie S1: Eps15 droplets fusion.** Time course of fusion events between condensates containing Eps15 droplets (green) in 20 mM Tris-HCl, 150 mM NaCl, 5 mM TCEP, 1 mM EDTA and 1 mM EGTA at pH 7.5 with 3 weight% PEG8000.

**Movie S2: Eps15 droplets fusion with the addition of Epsin1.** Time course of fusion events between condensates containing 16 μM Eps15 (green) and 1.6 μM Epsin1 (red). Condensates were incubated in 20 mM Tris-HCl, 150 mM NaCl, 5 mM TCEP, 1 mM EDTA and 1 mM EGTA at pH 7.5 with 3 weight% PEG8000.

**Movie S3: Eps15 droplets fusion with the addition of Epsin1 and K63 linkage TetraUb.** Time course of fusion events between Eps15 (16 μM) condensates (green) with 1.6 μM of Epsin1 (red) and 0.1 μM K63 linkage TetraUb (magenta), respectively. Condensates were incubated in 20 mM Tris-HCl, 150 mM NaCl, 5 mM TCEP, 1 mM EDTA and 1 mM EGTA at pH 7.5 with 3 weight% PEG8000.

